# Functional architecture of cerebral cortex during naturalistic movie-watching

**DOI:** 10.1101/2022.03.14.483878

**Authors:** Reza Rajimehr, Haoran Xu, Asa Farahani, Simon Kornblith, John Duncan, Robert Desimone

**Affiliations:** MRC Cognition and Brain Sciences Unit, University of Cambridge, Cambridge, UK; McGovern Institute for Brain Research, Massachusetts Institute of Technology (MIT), Cambridge, MA, USA; McConnell Brain Imaging Centre, Montreal Neurological Institute, McGill University, Montreal, Canada; Google Brain, Toronto, Canada; Department of Experimental Psychology, University of Oxford, Oxford, UK; Department of Brain and Cognitive Sciences, Massachusetts Institute of Technology (MIT), Cambridge, MA, USA

**Author notes:** Corresponding author:* Reza Rajimehr, MRC Cognition and Brain Sciences Unit, University of Cambridge, 15 Chaucer Road, Cambridge CB2 7EF, UK. These authors jointly supervised this work (John Duncan, Robert Desimone).

**Keywords:** cerebral cortex, cortical map, parcellation, clustering, fMRI, human connectome project

## Abstract

Characterizing the functional organization of cerebral cortex is a fundamental step in understanding how different kinds of information are processed in the brain. Neuroimaging studies over the past decades have uncovered the function of many cortical areas in response to selected stimulus categories. However, it is still unclear how cortical areas are organized during naturalistic visual and auditory stimulation. Here we used high-resolution functional MRI data from 176 human subjects to map the macro-architecture of the entire cerebral cortex based on responses to a 60-minute audiovisual movie stimulus. A data-driven clustering approach revealed a map of 24 functional areas/networks, each related to a specific aspect of sensory or cognitive processing. The map included three distinct executive control (domain-general) networks which showed a strong push-pull interaction with domain-specific regions in visual, auditory, and language cortex. The cortical parcellation scheme presented here provides a comprehensive and unified map of functionally defined areas, which could replace a large set of functional localizer maps.

## Introduction

Human cerebral cortex contains a mosaic of areas. These areas are typically delineated based on histology (cytoarchitecture and myeloarchitecture), topography, functional properties, and connectivity patterns (Felleman and Van Essen, 1991). In 1909, Korbinian Brodmann subdivided one cerebral hemisphere into 52 cytoarchitectonic areas (Brodmann, 1909). Recently Brodmann’s map has been refined through histological analysis of a large sample of brain tissues (Amunts et al., 2020). Modern neuroimaging techniques have also enabled cartographers to map topography, function, and connectivity of many cortical areas. Using multimodal neuroimaging data and a semi-automated gradient-based parcellation approach, Glasser et al. delineated 180 areas/parcels in each hemisphere of cerebral cortex (Glasser et al., 2016a).

In higher associative areas of the temporal and frontal lobes, the architectonic borders between areas are sometimes ambiguous due to gradual transitions in microstructure (Amunts and Zilles, 2015). These areas also show considerable variability in anatomical location relative to cortical folds (Fischl et al., 2008). In addition, topographic maps, which provide important landmarks for defining areas in early sensory and motor cortices (Wang et al., 2015b), are either absent or hard to resolve in these higher-tier areas. It appears that function and connectivity could be reliable features in partitioning the cortex when architectonic and topographic borders are not well-defined.

Functional connectivity analyses suggest that cortical areas are not isolated patches. Instead, each area is strongly coupled with a number of geographically distinct regions to form large-scale functional networks (Yeo et al., 2011; Ji et al., 2019). Arguably these macroscopic networks might be building blocks of cortical organization because each network is at least partially responsible for certain functions (e.g. a specific sensory processing and its cognitive modulations) which are ultimately relevant to behavior. Functional connectivity, as measured by fMRI, is usually based on correlating the activity time-courses during rest. Many functionally defined areas are not preferentially active in the absence of a stimulus, and therefore, it would be difficult to find fine-grained segregation between functional networks in the resting state. To identify such areas, a set of ‘functional localizer’ scans could be used, each for localizing a specific cortical area or a network of areas. These localizers have been very helpful in understanding the functional organization of high-level cortical areas (Kanwisher, 2017). However, designing localizer experiments for a large number of stimulus categories and task conditions would be inefficient and perhaps practically impossible.

Here we used rich audiovisual movie stimuli to effectively activate a large portion of cerebral cortex (sensory, category-selective, and cognitive regions). Then using a data-driven approach, the entire cortex was functionally parcellated based on similarity/commonality in the pattern of fMRI responses to the movie. The results revealed a comprehensive map of cortical areas, networks, and subnetworks during naturalistic movie watching. This parcellation reflects the architecture of cerebral cortex when it is involved in processing complex and dynamic audiovisual scenes.

## Results

We used movie-watching fMRI data of 176 healthy young adults from the Human Connectome Project (HCP) database (https://www.humanconnectome.org/study/hcp-young-adult). Subjects were scanned in a 7T scanner while watching short (ranging from 1 to 4.3 minutes in length) audiovisual movie clips. The clips were independent film and Hollywood movie excerpts which were concatenated and presented in four functional runs (total scan duration: 60 minutes) (**Figure 1a**). The movies contained a variety of visual stimuli (people, animals, scenes, and objects), visual actions, sounds, music, speech, linguistic and social communications, and sometimes narratives. There were also 20-second rest periods between the movies. Subjects were allowed to make free eye-movements during the scans. It has been shown that visual representations in high-level cortical areas are tolerant to eye movements when watching a natural movie (Nishimoto et al., 2017).

**Figure 1:**
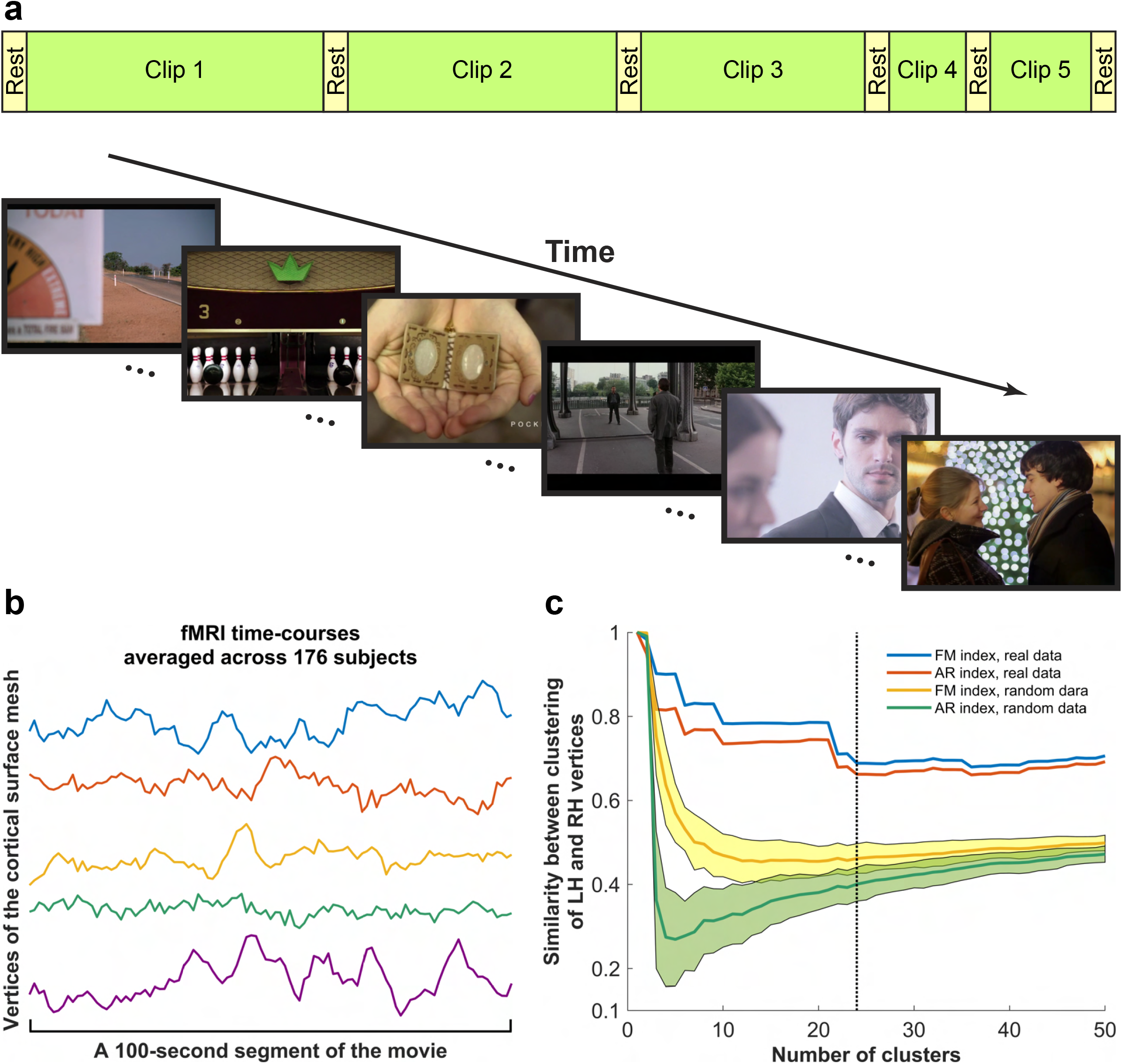
Naturalistic movie-watching paradigm and clustering analysis of fMRI data. (a) Subjects were scanned in a 7T scanner while watching audiovisual movie clips. In total, 18 clips were presented in four functional runs (5 clips in the first and third runs, 4 clips in the second and fourth runs). The last clip of the four runs was the same, and it was included for test-retest purposes. 20-second rest periods were interleaved between movie clips. (b) Examples of averaged fMRI time-courses. In each cortical vertex, time-courses were averaged across 176 subjects after de-meaning. (c) Clustering analysis was performed on vertices of the entire cortex, then similarity of clustering maps in the two hemispheres was computed for 50 levels of hierarchical clustering. Two different metrics, Fowlkes– Mallows (FM) index and adjusted Rand (AR) index, were used to quantify the clustering similarity. The clustering similarity was also computed for 100 permutations of simulated/random data. Before clustering, the random noise data were spatially smoothed on the surface using a Gaussian kernel with sigma = 4 mm, mimicking a hemodynamic point spread function of 4 mm (Engel et al., 1997). The shaded areas around the curves indicate one standard deviation, calculated based on 100 simulations. The dotted line indicates the level of 24 clusters.

Functional data in individual subjects were preprocessed and multimodally transformed to a standard cortical surface where left and right hemispheres were precisely registered to each other (i.e., there was a one-to-one correspondence between points/vertices of the two hemispheres) (Glasser et al., 2016b). Each hemisphere contained ~ 30,000 vertices. In each subject, time-courses of activity were de-meaned and concatenated across functional runs. Data matrices (vertices x time-points) were then averaged across subjects, assuming a robust inter-subject synchronization of cortical activity during natural vision (Hasson et al., 2004). Since the movies were presented once to the subjects, the inter-subject averaging of functional data provided more reliable activation patterns. Furthermore, idiosyncratic low-frequency fluctuations of fMRI response, which have an intrinsic origin, were largely subtracted out by inter-subject averaging (Kim et al., 2018), and the averaged time-courses closely reflected what was presented in the movies. Examples of averaged time-courses are shown in **Figure 1b**.

Next, we constructed an activity space in which each axis corresponded to the functional activity at a given time-point. Given the sampling rate of 1 Hz during data acquisition (TR = 1 sec), the averaged time-courses included 3655 time-points for the entire scan session. Thus, the activity space contained 3655 orthogonal axes. Vertices of the two hemispheres (~ 60,000 vertices) were data-points in this space. Our primary goal was to find distinct clusters of vertices based on the geometric distance between data-points in the activity space. For the clustering analysis, a hierarchical clustering algorithm was used. Unlike other clustering algorithms (such as *k*-means clustering) in which the number of clusters is fixed and arbitrarily predefined, the hierarchical clustering groups data-points at various levels/scales. This multi-scale approach can be particularly useful for testing hierarchical (‘coarse-to-fine’) partitioning of the spatially organized maps. At each level of clustering, a color was assigned to vertices within each cluster, then the colored vertices were visualized on 2-D flat patches of cortex (**Supplementary Figure 1**). Although we did not include any information about the location of vertices in the clustering analysis, the maps demonstrated a remarkable spatial organization of functionally defined clusters on the cortical surface. At the very top level of hierarchical clustering, the first cluster appeared in visual cortex of the occipital lobe. By progressively increasing the number of clusters, visual cortex and the remaining parts of cerebral cortex were recursively subdivided into smaller clusters, and the resulting maps showed macro-organization of cerebral cortex at finer scales.

To assess the reliability of the clustering maps, we computed the similarity between cluster labels of vertices in the two hemispheres (note that each vertex in the left hemisphere had a corresponding vertex in the right hemisphere). Using two different metrics, Fowlkes–Mallows index (Fowlkes and Mallows, 1983) and adjusted Rand index (Hubert and Arabie, 1985), we observed a high degree of similarity between clustering of vertices in the two hemispheres (**Figure 1c**). In a control analysis, the clustering similarity between the two hemispheres was computed for simulated data in which the real functional activities of vertices were replaced with the Gaussian white noise. The clustering similarity was significantly higher for real data, compared to simulated/random data (for every level of clustering: Bonferroni-corrected p < 0.01, permutation test) (**Figure 1c**). For real data, the clustering similarity gradually decreased as the number of clusters increased, but the similarity values remained somewhat stable after the level of 24 clusters. We used 24 as a cutoff point for the hierarchical clustering to investigate the functional parcellation of cerebral cortex.

**Figure 2** shows the functional parcellation maps at the level of 24 clusters. The maps in left and right hemispheres were largely similar. To gain a better insight about the anatomical location of clusters, a multimodal parcellation of cerebral cortex (a parcellation with 180 cortical areas in each hemisphere) (Glasser et al., 2016a) was overlaid on our maps. Clusters could be classified into four groups: 1) clusters within sensory cortices, 2) clusters corresponding to the category-selective areas, 3) clusters corresponding to major cognitive networks, and 4) a cluster corresponding to anterior temporal cortex and other cortical regions with low signal-to-noise ratio in fMRI.

**Figure 2:**
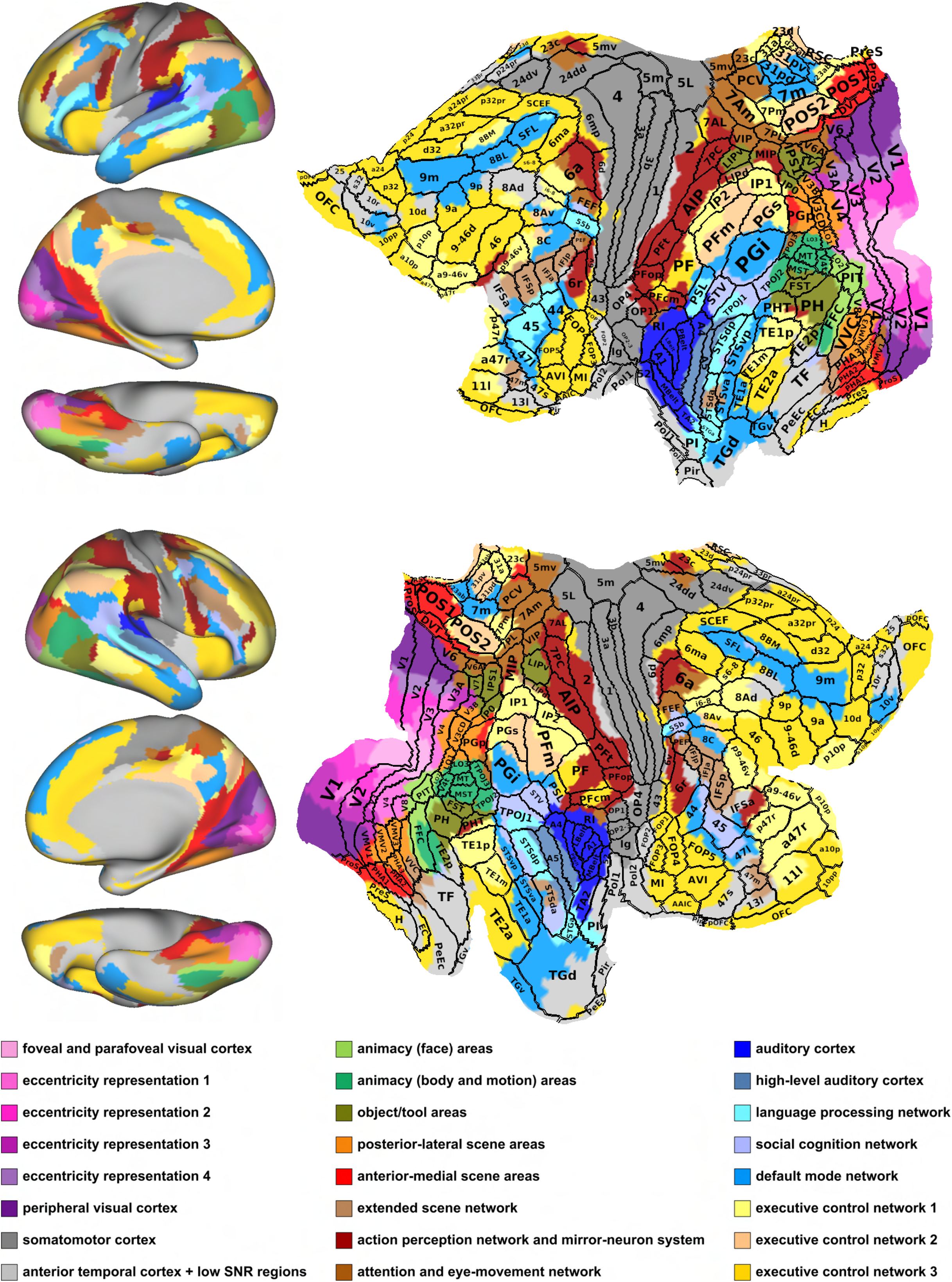
Parcellation maps at the level of 24 clusters in the hierarchical clustering analysis. On the left, maps are displayed on lateral, medial, and ventral views of the inflated cortical surface (fs_lr surface). Vertices of the medial wall were not included in the analysis. On the right and in the subsequent figures, maps are displayed on 2-D flat patches of fs_lr surface so that the entire cortex could be seen in a single view. The top and bottom rows show parcellation maps in left and right hemispheres, respectively. Borders and areal names of a multimodal parcellation (an atlas of 180 cortical areas in each hemisphere) (Glasser et al., 2016a) were overlaid on our parcellation maps. The clusters were named based on their anatomical location and topographic correspondence with previously described functional areas/networks in cortex.

Six clusters in early visual cortex (V1-V4) were arranged along dorsal-ventral axis of the occipital lobe, and they appeared to correspond to the representations of visual field eccentricities, from foveal to peripheral visual fields. Two clusters corresponded to auditory cortex (A1, belt, and parabelt) and high-level auditory cortex (A4 and A5). One large cluster contained several areas in somatomotor cortex. Seven category-selective clusters included animacy (face) areas, animacy (body and motion) areas, object/tool areas, posterior-lateral scene areas, anterior-medial scene areas, extended scene network, and action perception network (aka ‘mirror-neuron’ system). In **Figures 4,6**, we will comprehensively evaluate the correspondence between these clusters and category-selective areas localized through conventional localizer maps. Seven cognitive processing clusters included attention and eye-movement network, language processing network, social cognition network, default mode network, and three executive control networks. In **Figures 5-7**, we will comprehensively evaluate the correspondence between these clusters and functional networks identified through other analyses.

**Figure 3:**
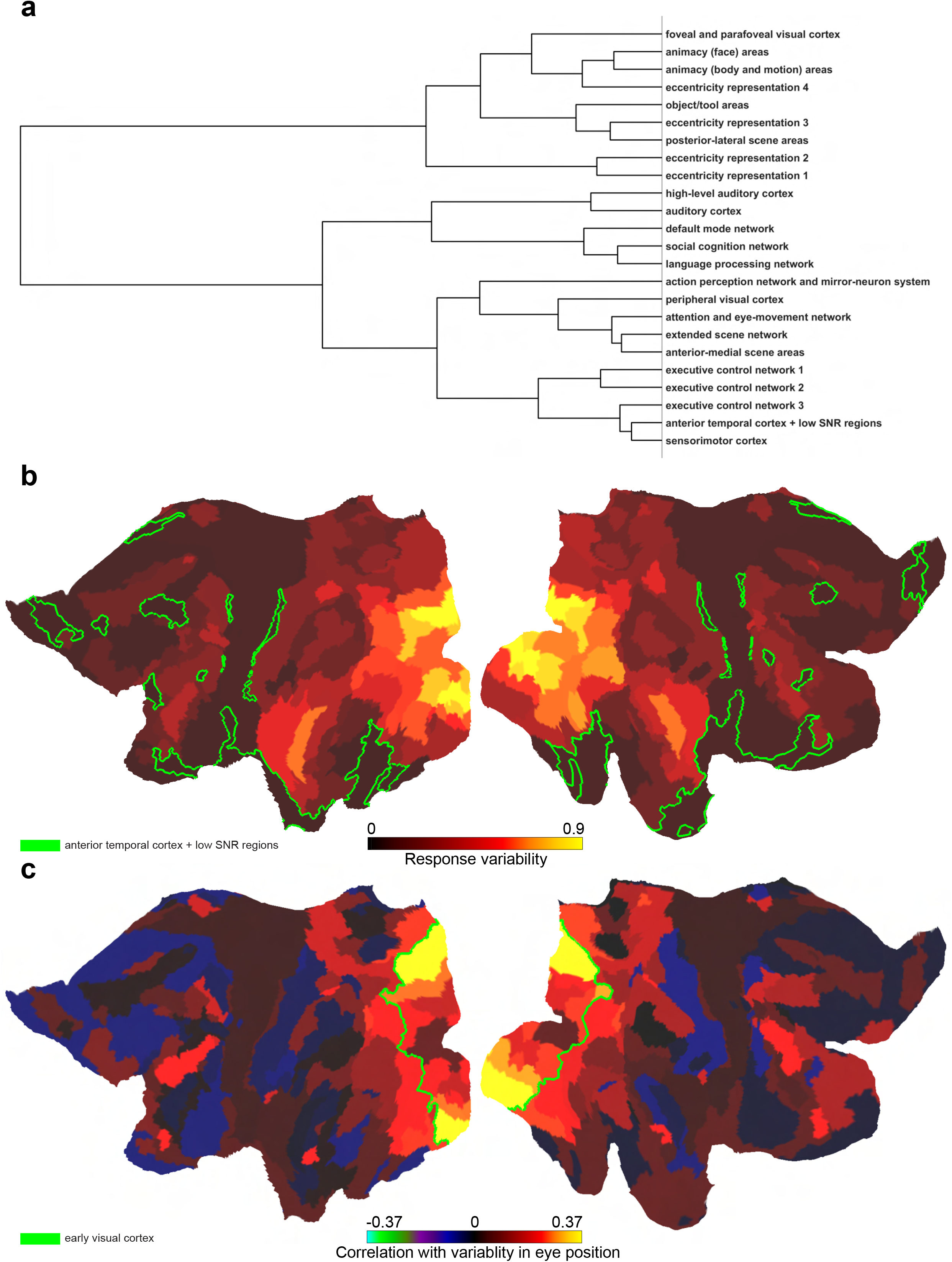
Hierarchical clustering tree and basic response properties of the clusters. (a) Dendrogram of hierarchical clustering tree from 2 to 24 clusters. (b) The maps show response variability in 24 clusters. Response variability was estimated by calculating the standard deviation of the mean time-course of activity in each cluster. Regions indicated by green borders, somatomotor cortex, and executive control network 3 showed low response variability. (c) The maps show the correlation between the mean time-course of activity in each cluster and the variability in eye position. Before computing the correlation, the activation vectors were shifted by four seconds to account for the hemodynamic response delay (Menon et al., 1995). For every second of the movie-watching scan, the variability in eye position in each subject was defined as the square root of sum of variances of horizontal and vertical eye position. If a time-point contained blinks or an abrupt eye-tracking signal loss, a NaN value was assigned to that time-point. Thus, for 3655 time-points of the movie-watching scan, we obtained values indicating the amount of eye-movements. The values were averaged across subjects, then the averaged vector was correlated with the activation vectors in 24 clusters. Early visual cortex included six clusters in foveal, mid-peripheral, and peripheral visual cortex.

**Figure 4:**
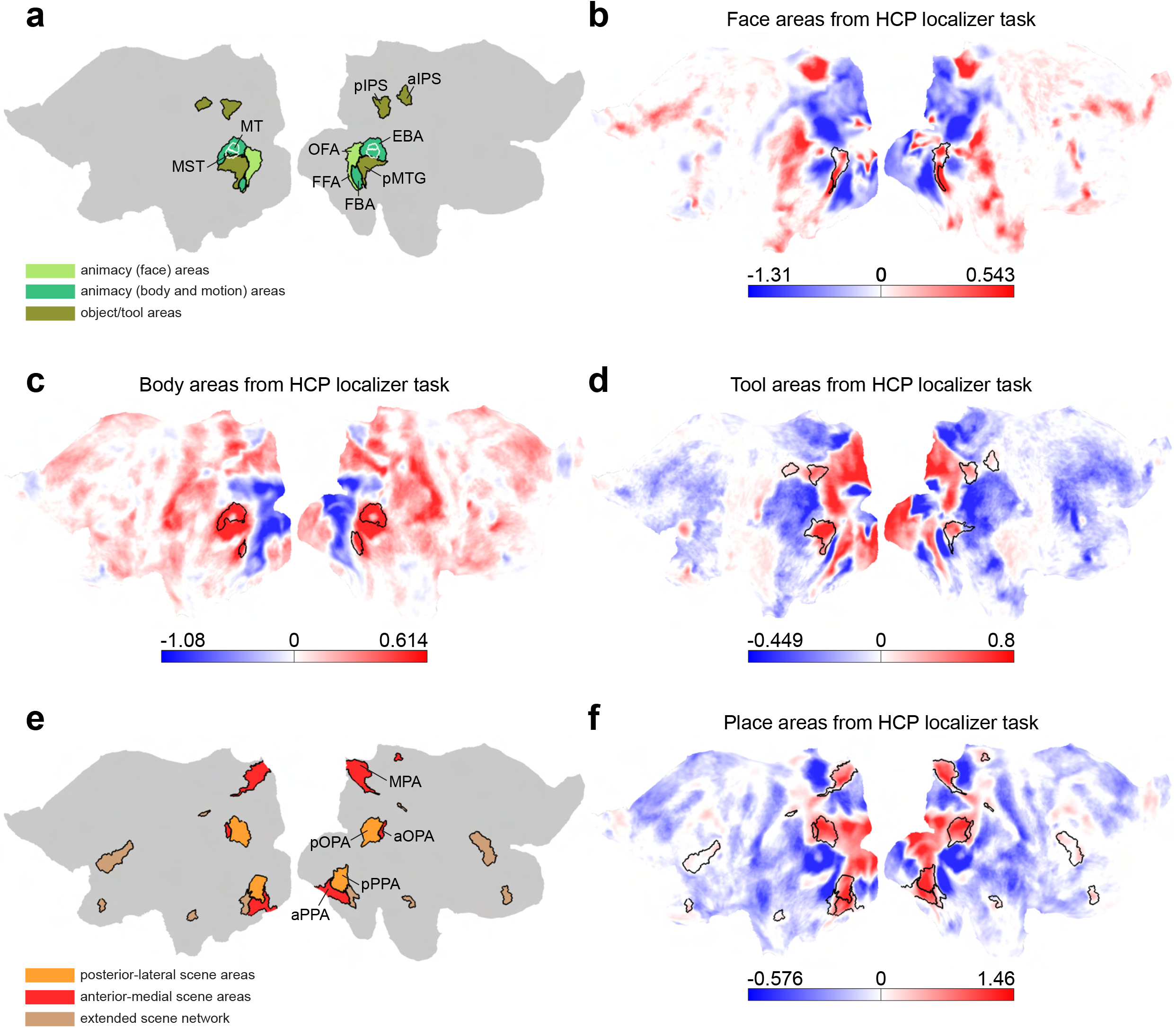
Topographic relationship between category-selective clusters and functional localizer maps. The animacy and object/tool clusters are shown in panel a, and the scene clusters are shown in panel e. The maps in panels b-d and f are group-average functional localizer maps for visual categories (faces, bodies, tools, and places) from S1200 package, and they were obtained from the HCP working memory task by comparing the activation for one category versus the average activation for the other three categories. The maps represent Cohen’s d effect size, and borders of relevant clusters are shown on the maps. Using localizer maps as a guide, different subparts of the clusters were found to be analogous to previously described category-selective areas (see text for a full name of the areas). The area MT/MST (white outlines) was derived from the multimodal cortical parcellation.

**Figure 5:**
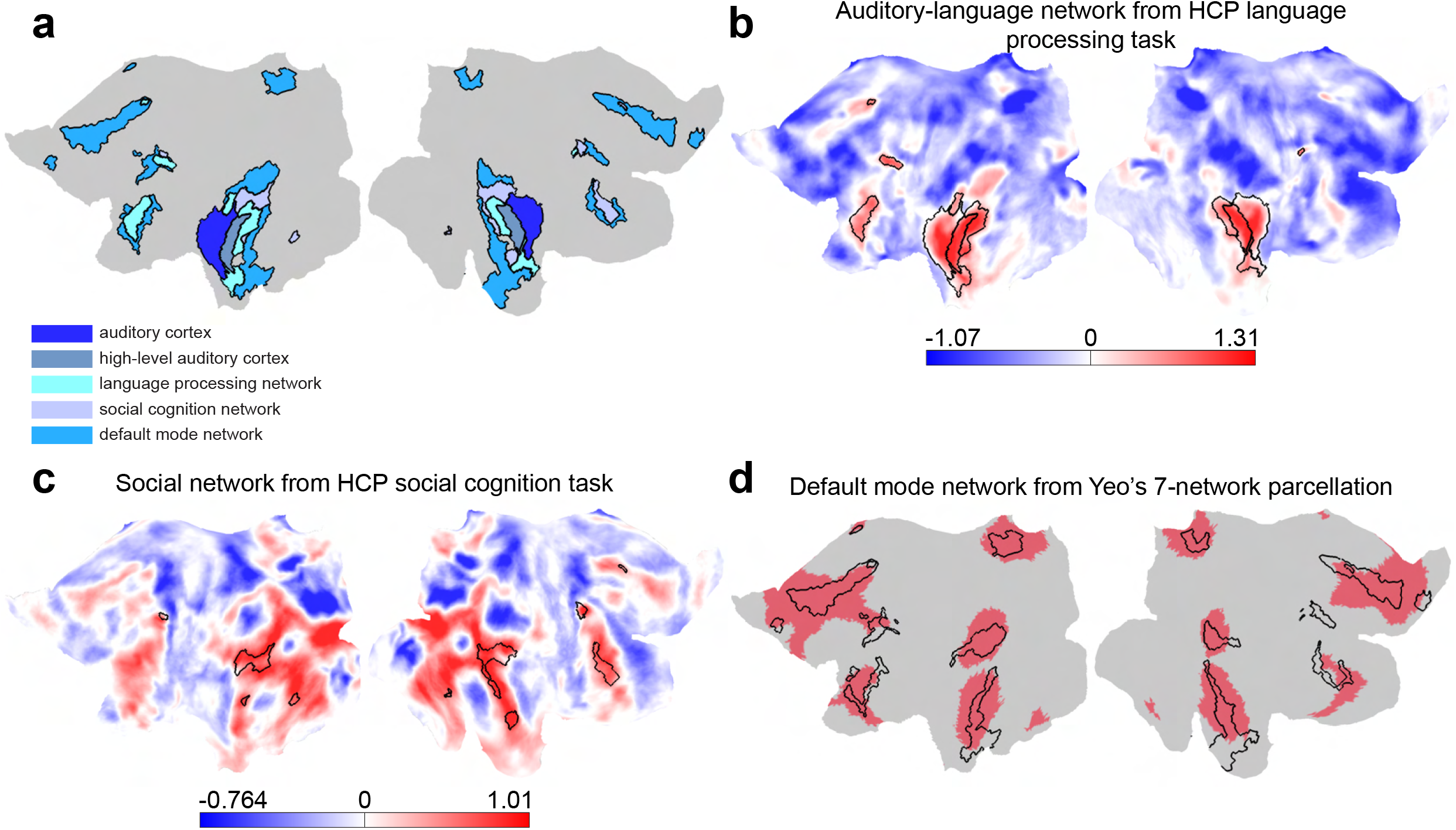
Topographic relationship between lateral temporal clusters and some cognitive networks. (a) Five clusters arranged dorsoventrally in lateral temporal cortex. (b) Group-average activation map from S1200 package for the contrast of stories vs. baseline in the HCP language processing task. The baseline condition in this task represented the mean activity across all time-points in each run. (c) Group-average activation map from S1200 package for the contrast of social vs. random stimuli in the HCP social cognition task. In the social cognition task, subjects were presented with short video clips of simple geometric shapes (squares, circles, triangles) either interacting in some way (Social condition) or moving randomly (Random condition). The maps in panels b-c represent Cohen’s d effect size. (d) Default mode network from Yeo’s 7-network parcellation. In panels b-d, borders of relevant clusters are shown.

**Figure 6:**
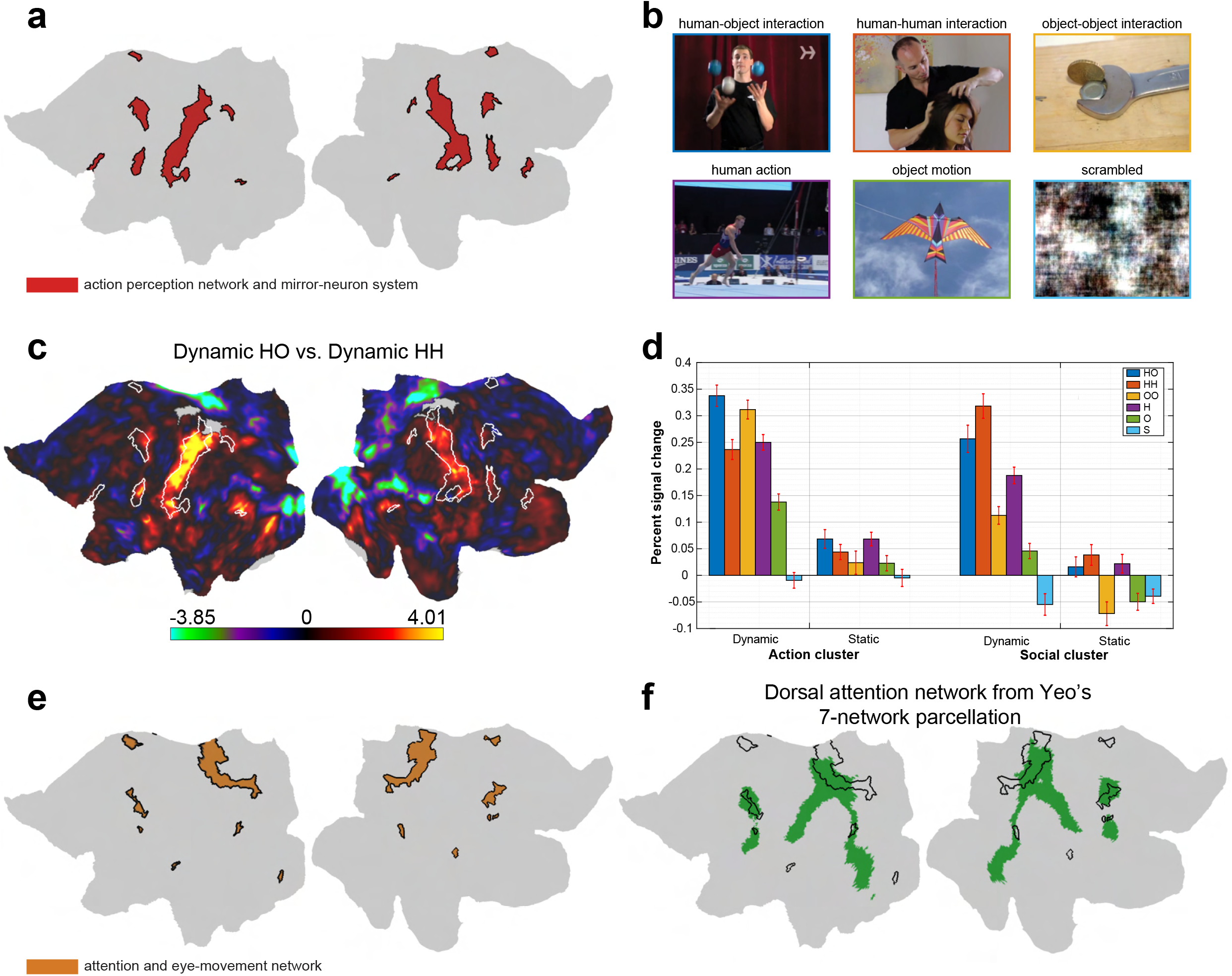
Characterization of two clusters located in parietal cortex. (a) A cluster labeled as action perception network and mirror-neuron system. (b) The action categories used in the action localizer experiment. (c) Mixed-effects group-average maps for the contrast of dynamic human-object interactions (yellow activations) vs. dynamic human-human interactions (cyan activations) based on fMRI data from an independent group of 22 subjects. Data were analyzed in FreeSurfer on the fsaverage surface (see Methods for more details), then the activation maps were resampled onto the fs_lr surface using spherical transformation. The maps show FDR-adjusted significance values in a logarithmic format. (d) The bar plot shows the percent signal change values for dynamic and static stimuli of six action categories in the action perception and social cognition clusters. The percent signal change values were computed based on the contrast of each stimulus condition vs. fixation. For the social cognition cluster, only vertices of the right hemisphere were included in the analysis due to a strong hemispheric lateralization of this cluster. Error bars indicate one standard error of the mean across subjects. (e) A cluster labeled as attention and eye-movement network. (f) Dorsal attention network from Yeo’s 7-network parcellation. In panels c and f, borders of relevant clusters are shown.

**Figure 7:**
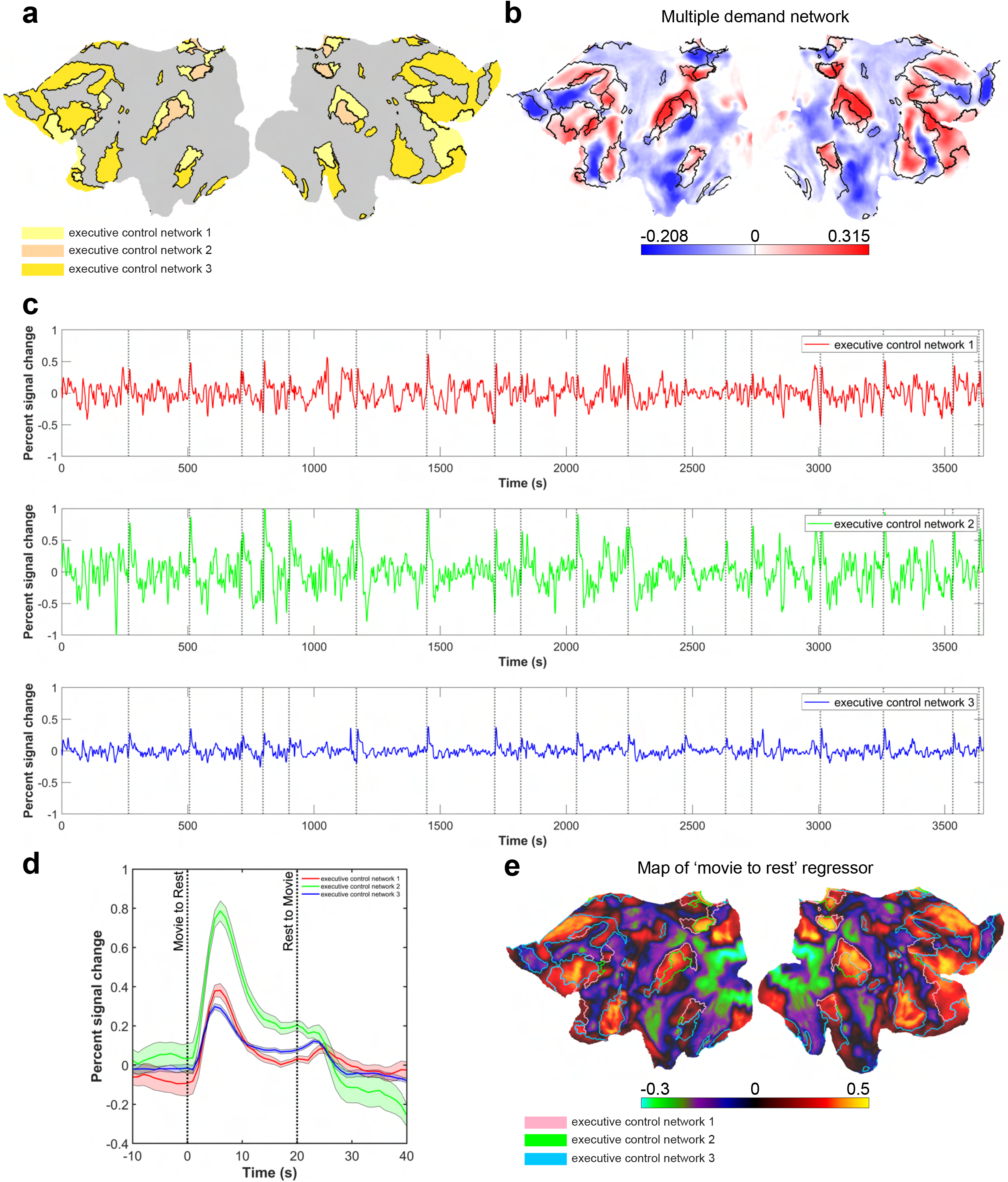
Three executive control clusters and their functional properties. (a) Three clusters labeled as executive control network 1, 2, 3. (b) The multiple demand network identified using the task fMRI data of 449 HCP subjects. The maps represent the averaged percent signal change values. (c) The mean time-course of activity in the executive control networks. The dotted lines indicate the onset of 20-second rest periods. (d) The graph shows the averaged responses across selected time-windows around 20-second rest periods. The shaded areas indicate one standard error of the mean across 18 time-windows. (e) A regressor was defined based on the times of transition from movie to rest. The regressor was convolved with a canonical hemodynamic response function, then it was correlated with the time-courses of all cortical vertices. In panels b and e, borders of executive control clusters are shown.

**Figure 8:**
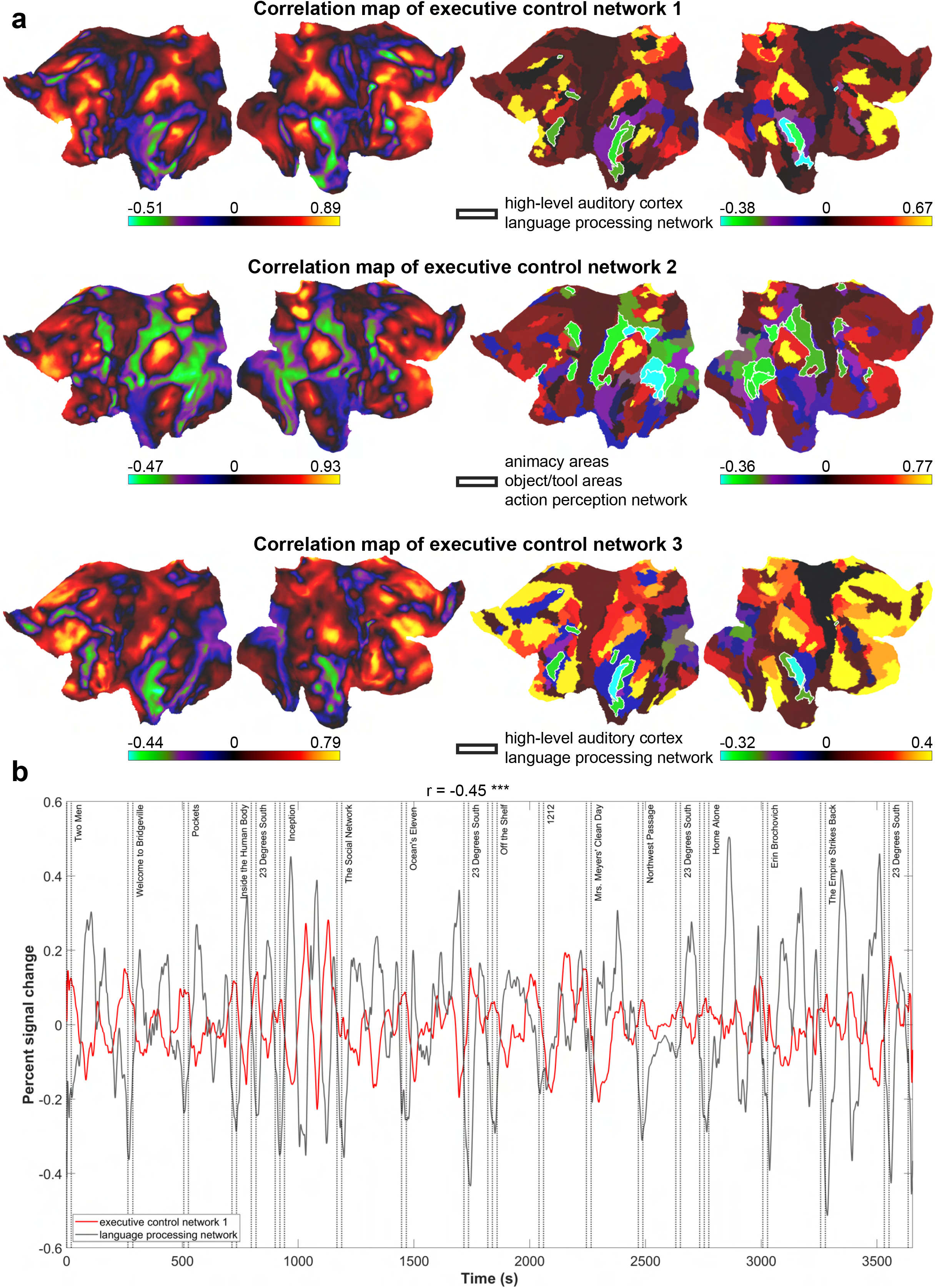
A push-pull interaction between executive control networks and domain-specific areas of cerebral cortex. (a) The maps show the correlation between the mean time-course of activity in the executive control clusters and the time-courses of activity in all cortical vertices (the maps on the left) and cortical clusters (the maps on the right) after removing 20-second rest and 20-second after-rest periods. Domain-specific clusters are highlighted on the cluster-wise maps. (b) As an example, the mean time-courses of activity in two clusters are demonstrated. The time-courses were smoothed with a moving average window of 50 seconds, just for visualization purposes (the correlation analysis was done without smoothing of the data). The onset of all movie clips is indicated in the plot. The correlation coefficient value was computed after removing 20-second rest and 20-second after-rest periods. ***: p << 0.0005.

The hierarchical clustering tree is shown in **Figure 3a**. Tracing of clusters in the clustering tree revealed some interesting features. One branch of the tree included foveal/parafoveal visual cortex and animacy areas, whereas another branch included peripheral visual cortex and scene areas. Such a distinction is consistent with some current models for the origin of animacy/face and scene selectivity in visual cortex (Levy et al., 2001; Hasson et al., 2003; Arcaro and Livingstone, 2017). Moreover, default mode network, social cognition network, and language processing network were derived from a common node in the tree structure, suggesting a link between semantic, social, and linguistic representations in cerebral cortex (Rajimehr et al., 2021).

We investigated two basic response properties in all clusters. In one analysis, we looked at the response variability/fluctuation of the mean time-course in each cluster. The result of this analysis would indicate how effectively the clusters were activated-deactivated by the movie stimulus. As shown in the maps in **Figure 3b**, a cluster that included anterior temporal cortex and some scattered patches in the frontal lobe showed the lowest response variability. This cluster was located in cortical regions where the fMRI signal-to-noise ratio is typically low due to susceptibility artifacts (Rajimehr et al., 2009). In another analysis, we looked at the correlation between the mean time-course of activity in each cluster and the variability in eye position. Variability in eye position was calculated by analyzing behavioral eye-tracking data from all subjects. As shown in the maps in **Figure 3c**, there was a systematic increase in correlation from foveal to peripheral regions in early visual cortex, consistent with the fact that subjects made more eye-movements when there were more items in the peripheral visual field.

To examine the topographic relationship between some of the clusters in occipito-temporal cortex and classically defined category-selective areas, we qualitatively compared these clusters with functional localizer maps for visual categories (**Figure 4**). These group-average maps of ~ 1000 HCP subjects were obtained by comparing blocks of one category versus blocks of other categories (e.g. faces vs. bodies, tools, and scenes). The animacy (face) cluster showed overlap with face-selective vertices in regions we identified as occipital face area (OFA) and fusiform face area (FFA) (Kanwisher and Yovel, 2006) (**Figure 4a,b**). The animacy (body and motion) cluster showed overlap with motion-selective vertices in MT/MST and body-selective vertices in regions we identified as extrastriate body area (EBA) and fusiform body area (FBA) (Peelen and Downing, 2005) (**Figure 4a,c**). The object/tool cluster showed overlap with tool-selective vertices in posterior middle temporal gyrus (pMTG), posterior intraparietal sulcus (pIPS), and anterior intraparietal sulcus (aIPS) (Beauchamp et al., 2003) (**Figure 4a,d**). Two scene clusters showed overlap with scene-selective vertices in regions we identified as occipital place area (OPA), parahippocampal place area (PPA), and medial place area (MPA) (Epstein and Baker, 2019) (**Figure 4e,f**). Specifically, one cluster included posterior OPA and posterior PPA, and the other cluster included anterior OPA, anterior PPA, and MPA. This functional segregation within scene processing network has also been reported previously (Baldassano et al., 2013; Nasr et al., 2013). For scenes, tools, and bodies, each cluster contained patches separated by a relatively large distance on the cortical surface. This property, which was seen in many clusters of our parcellation map, suggests that clusters have not resulted from artifactual correlations (spatial autocorrelations (Kriegeskorte et al., 2008)) between the fMRI hemodynamic responses of neighboring voxels.

The results above were quantitatively confirmed in a region-of-interest (ROI) analysis. In each category localizer map, the corresponding clusters from the movie data showed the highest activation (**Supplementary Figure 2a**). Furthermore, the face, body, object/tool, and scene clusters were generally more active/responsive for frames of the movie that included categories related to those clusters (faces and people in the face cluster; body parts and hands in the body cluster; objects, tools, texts, and eyes in the object/tool cluster; indoor and outdoor scenes in the scene clusters) (**Supplementary Figure 2b**). One cluster, named extended scene network here (**Figure 4e**), also showed a relatively high response to scenes compared to most other categories (**Supplementary Figure 2a**), and its preferred movie frames included pictures of scenes (**Supplementary Figure 2b**). This cluster was closely related to the anterior-medial scene cluster in the hierarchical clustering tree (**Figure 3a**), had a large component in lateral prefrontal cortex, and might be involved in processing high-level semantic aspects of scenes in a naturalistic condition (Popham et al., 2021).

Five clusters were arranged dorsoventrally in lateral temporal cortex. These clusters were named auditory cortex, high-level auditory cortex, language processing network, social cognition network, and default mode network (**Figure 5a**). The lateral temporal component of the language processing cluster was located in dorsal superior temporal sulcus (STSd) and area PSL of the left hemisphere, with a smaller accompanying component in the right hemisphere. Additional components of this cluster were located in lateral prefrontal cortex of the left hemisphere, within areas SFL, 55b, and Broca’s area (Brodmann areas 44, 45, and 47). The auditory and language clusters matched almost perfectly with the activations produced by the HCP language processing task (the comparison between auditorily presented stories versus baseline) (**Figure 5b**). The main component of the social cognition cluster was located at the temporo-parietal junction (areas TPOJ1 and STV). Interestingly, smaller components of this cluster were located in specific areas in lateral temporal and lateral prefrontal cortex of the right hemisphere, which were homotopic (corresponding) to the left-hemisphere language areas. This feature of our parcellation map was consistent with a recent demonstration of complementary hemispheric lateralization of language and social processing in the human brain (Rajimehr et al., 2021). The social cluster was located within the areas activated by the HCP social cognition task (the comparison between visually presented social versus random stimuli) (**Figure 5c**). A more extended activation pattern produced by this task could be due to uncontrolled visual confounds. The default mode cluster was located in ventral STS and temporal pole (area TG). The non-temporal components of this cluster were located in area PGi, medial parietal cortex, medial prefrontal cortex, and regions near/surrounding 55b and Broca’s area. All these components overlapped with the default mode network obtained from a functional connectivity analysis of resting-state fMRI data (Yeo et al., 2011) (**Figure 5d**).

An additional category-selective cluster, located in lateral parietal and premotor cortex, appeared to correspond to the action perception network (mirror-neuron system) (Buccino et al., 2001) (**Figure 6a**). We confirmed this by analyzing data from an independent block-design fMRI experiment in which dynamic videos and static images from six action categories were presented to 22 subjects (**Figure 6b** and **Supplementary Figure 3**). The action categories included human-object interaction (HO), human-human interaction (HH), object-object interaction (OO), human action (H), object motion (O), and scrambled condition (S). Group-average maps of univariate comparison between dynamic HO vs. dynamic HH revealed a localized activation pattern for HO, which matched almost perfectly with the action perception cluster (**Figure 6c**). This result suggests that the action perception network in parietal and premotor cortex responds preferentially to specific types of action stimuli which involve human-object interactions. The HO activations were stronger in the left hemisphere. In an ROI analysis (**Figure 6d**), the action perception cluster showed significantly higher response to dynamic HO compared to other conditions (p < 0.05 for all pairwise comparisons between dynamic HO and other conditions, except dynamic HO vs. dynamic OO; repeated-measures ANOVA, Tukey post-hoc test). In contrast, the social cognition cluster in the right hemisphere showed significantly higher response to dynamic HH compared to other conditions (p < 0.05 for all pairwise comparisons between dynamic HH and other conditions; repeated-measures ANOVA, Tukey post-hoc test).

Adjacent to the action perception cluster, there was a cluster which overlapped with frontal eye field (FEF) and superior parietal parcels of the multimodal cortical parcellation (**Figure 2** and **Figure 6e**). A meta-analysis of fMRI activations has demonstrated that FEF and regions in/near the intraparietal sulcus are consistently activated during attention tasks (He et al., 2007). Thus, based on the anatomical location, we predicted that this cluster is related to the attention and eye-movement network. To test this prediction, we evaluated the topographic correspondence between this cluster and the dorsal attention network (Corbetta and Shulman, 2011). The dorsal attention network was identified through a functional connectivity analysis of resting-state fMRI data (Yeo et al., 2011). As shown in **Figure 6f**, the attention and eye-movement cluster showed partial overlap with this network. A possible reason for partial overlap is that some dorsal attention regions outside the cluster (e.g. MT/MST and action perception areas) were strongly activated by other components of the movie, and therefore they were assigned to other networks.

A large swath of parietal, temporal, and prefrontal cortex was occupied by three clusters which were named executive control network 1, 2, 3 (**Figure 7a**) based on the anatomical location and also a large overlap with the multiple demand network (**Figure 7b**). The three clusters were spatially juxtaposed throughout the cortex. The multiple demand network was identified by averaging the HCP group-average beta maps of three task contrasts (2-back vs. 0-back working memory task, hard vs. easy relational processing task, and math vs. story task) (Assem et al., 2020). The multiple demand or domain-general network is believed to be flexibly involved in the execution of many tasks, and it may play a core role in cognitive control (Duncan, 2010). To explore the role of executive control clusters during passive movie-watching paradigm, we first looked at the mean time-course of activity across all cortical vertices within these clusters. The time-courses revealed a surprisingly large response at the transition from movie to rest periods (**Figure 7c**). This large and significant response was quite evident, especially in executive control network 2, when the responses were averaged across selected time-windows around 20-second rest periods (**Figure 7d**). Such transient response was either weak or absent at the transition from rest to movie periods. To evaluate this transient response in the entire cortex, we defined a regressor based on the times of transition from movie to rest, then computed the correlation between the regressor and the time-courses of all cortical vertices (**Figure 7e**). Regions of high positive correlation were localized in executive control networks. As expected, the stimulus-driven regions in early visual and auditory cortex showed strong negative correlation, though intriguingly, an additional region of the most peripheral visual cortex showed the same positive response seen in executive control networks.

Next, we measured the functional correlation between the mean time-course of activity in each executive control cluster and the time-courses of activity in all cortical vertices and cortical clusters after removing 20-second rest and 20-second after-rest periods. The vertex-wise and cluster-wise correlation maps are shown in **Figure 8a**. The three executive control networks were similar in their positive correlations, covering the whole multiple demand areas. Interestingly, the maps also revealed strong negative correlation between executive control networks and cortical regions/clusters which corresponded to domain-specific areas. Executive control network 1 and 3 had strong negative correlation with high-level auditory cortex and language processing network. Executive control network 2, which was located in areas POS2 and PFm (**Figure 2**), had strong negative correlation with high-level visual cortex including animacy areas, object/tool areas, and action perception network. Such anti-correlations were evident throughout the movies, for all the movie clips (**Figure 8b**). These results suggest a ‘push-pull’ interaction between domain-general and domain-specific areas of cortex. We did not observe regions of high negative correlation when the correlation maps were obtained using the HCP resting-state fMRI data (**Supplementary Figure 4**), suggesting that the push-pull interaction was driven by movie events.

## Discussion

In this human fMRI study, we used a data-driven approach to functionally parcellate the entire cerebral cortex. In this approach, we used rich audiovisual movie stimuli to drive the cortex and elicit a large variation in the patterns of response across voxels/vertices. In each vertex, the time-courses of response were averaged across subjects, considering that the local fMRI responses show remarkable inter-subject synchrony under natural viewing conditions (Hasson et al., 2004). The averaging was done in a common anatomical space after multimodal transformation of individual subjects’ data. Other methods of averaging (such as ‘hyperalignment’ (Haxby et al., 2011)) may improve the estimation of group-average fMRI responses. In the next step, we applied a clustering algorithm on data-points (vertices) in the activity space. We used hierarchical clustering which had an advantage of defining clusters/parcels at different scales/resolutions. The parcellation map at the level of 24 clusters showed clusters which topographically corresponded to previously known cortical areas and networks (e.g. the category-selective areas). This map is suitable for evaluating large-scale cortical networks (‘supra-areal organization’) (Buckner and Yeo, 2014). A parcellation map with a higher number of clusters could reveal finer distinctions within the networks and perhaps even millimeter-scale subregions within the areas. For example, some of the clusters in temporal and prefrontal cortex may contain a fine-grained representation of semantic information (Huth et al., 2016).

Previous studies have used resting-state fMRI data to parcellate cortex in humans. Yeo et al. identified a set of cortical networks using a clustering analysis applied on data from a population of 1,000 healthy subjects (Yeo et al., 2011). Other studies employed various computational techniques to parcellate individual subjects’ cortices (e.g. Wig et al., 2014; Laumann et al., 2015; Wang et al., 2015a). Using boundary mapping (Cohen et al., 2008) and graph theory, Nelson et al. parcellated the lateral parietal cortex into six distinct ‘modules’ (Nelson et al., 2010). The boundary mapping technique has been also used to parcellate the whole cortex (Gordon et al., 2016). In a follow-up study, a whole-cortex parcellation was obtained by integrating local gradient and global similarity approaches (Schaefer et al., 2018). Using a module detection algorithm, Goulas et al. parcellated the lateral frontal cortex (Goulas et al., 2012). Using *k*-means clustering, Kahnt et al. parcellated the orbitofrontal cortex (Kahnt et al., 2012). Finally, in one fMRI study (Vul et al., 2012), human subjects were scanned while viewing rapid event-related presentations of 69 unique images drawn from 9 object categories. Using data-driven clustering of voxels, this study found face, place, and body clusters/systems in the ventral visual pathway.

Some of the clusters in our parcellation map roughly correspond to the equivalent clusters in other parcellation maps that are based on resting-state data (e.g. default mode network in our parcellation and Yeo’s parcellation). However, the exact topography of clusters varies depending on the type of parcellation. In addition, the category-selective clusters, which are well differentiated in our map, are not clearly identified when the resting-state data are used. As mentioned in the Introduction, these clusters/regions of cortex are not preferentially active in the absence of a stimulus. Another advantage of using movie-watching data for parcellation is that, by analyzing the movie content, one can assess functional selectivity in less-studied cortical regions (i.e., regions for which a good a priori hypothesis about their function does not exist). These regions may have a counterintuitive selectivity to complex stimuli or a combination of stimuli. Such selectivity could be discovered through data-driven approaches.

Functional parcellation approaches have several advantages over classical localizer experiments in defining cortical areas. First, areas defined by a parcellation approach would have well-constrained selectivity to a specific stimulus due to the rich content of movie stimuli. Localizer experiments are typically designed for testing responses to a limited set of stimuli, and localizer maps sometimes show a widespread activation pattern. Some weak activations in these maps may actually disappear if the responses are tested for a broader range of stimuli. Second, a functional parcellation map may reveal genuine topographic borders between areas because it is based on responses obtained during a naturalistic condition where many cognitive processes and top-down modulations are present. Third, based on a parcellation/clustering map, one can explore the response properties in areas that have not been charted previously. For each cluster, one can use a ‘reverse-correlation’ approach to find the movie frames or the movie segments that produce the highest (and the lowest) response. These movie frames/segments can be used as a guide to deliver a set of hypotheses about the function of these clusters. These hypotheses and predictions could be thoroughly tested in well-controlled experiments. Fourth, the parcellation data can be used to investigate the functional interactions between different areas/networks of cortex during movie-watching. We took such an approach in **Figure 8** to look at the correlation maps of executive control networks.

Linking the fMRI responses to the movie frames can be useful for addressing further interesting questions. The parcellation algorithm forces a set of data-points to be segmented into discrete clusters with ‘hard borders’ between them. However, functional selectivity may not be homogeneous within a cluster. Instead, selectivity may change smoothly within and across the clusters. To clarify whether such smooth transitions exist, one can investigate the preferred movie frames for a region of cortex located at/near the border between two clusters. Furthermore, by comparing the preferred movie frames across clusters, subtle variations in response profile might be identified for clusters that show a common preference for a stimulus category. For instance, there might be a systematic difference between the preferred scene images of posterior-lateral versus anterior-medial scene clusters.

A movie sometimes has highly correlated information. For instance, faces and bodies are normally present together in the same frames of the movie. There is also co-occurrence of faces/bodies and motion in many frames of the movie since faces and bodies, as animate objects, are typically in motion. These stimulus correlations, which are also present during natural vision in everyday life, may have fundamental consequences on the cortical representation of these stimuli. First, stimuli that normally co-occur in natural vision may be represented in cortically adjacent regions. This is in fact the case for face, body and motion areas. Second, these areas may be partially activated by non-preferred but closely related stimuli. Accordingly, recent evidence suggests that biological movements contribute strongly to the responses in macaque face patches during the free viewing of movie clips (Russ and Leopold, 2015). These partial activations plus co-occurrence of stimuli in the movie would make it difficult to separate the corresponding areas by a clustering algorithm. However, in the parcellation map of 24 clusters, we were able to separate the face and body/motion clusters. The movie clips used in our study had a rich content, occasionally including close-up views of faces or body parts / hands. These particular movie frames may have helped to separate the face and body/motion clusters by producing a relatively higher response in one cluster or another. Body and motion areas, though grouped together in the map of 24 clusters, were separated at a later stage of hierarchical clustering (**Supplementary Figure 5**). Again parts of the movie, which include static people/bodies or dynamic inanimate objects, may have helped to distinguish these areas.

The cortical parcellation map in our study was based on group-average time-courses. Due to the lack of stimulus repetitions in the movie, the statistical power was inherently low. Therefore, extensive signal-averaging across subjects enabled us to have a better estimate of fMRI responses. The clusters obtained with this parcellation could be considered the most robust clusters that presumably play key roles in cortical processing. However, it is possible that each subject has an idiosyncratic parcellation map. By scanning individual subjects in multiple sessions, one can assess individual differences in the parcellation maps and in the layout of cortical areas/networks. Single-subject parcellation could also reveal small areas in ‘balkanized’ regions of cortex (e.g. face patches in anterior temporal cortex). These areas tend to be lost during the averaging process due to a high degree of variability in their locations. In classical localizer experiments, the size and topography of cortical areas in an individual subject depends on the amount of thresholding in the activation maps. This arbitrariness in threshold setting, which is a serious problem when comparing areas across subjects, can be avoided by parcellating the cerebral cortex using a clustering algorithm.

The parcellation map showed clusters in early visual cortex that spatially corresponded to the representation of eccentricity bands. This finding is consistent with the idea that ‘eccentricity bias’ is the major organizing principle in the visual occipito-temporal cortex (Malach et al., 2002; Hasson et al., 2002; Hasson et al., 2003). Interestingly, widespread correlation patterns of resting-state fMRI signal across early visual cortex also reflect topographic (eccentricity-based) organization (Arcaro et al., 2015). Thus, shared eccentricity representations may outweigh functional differences across anatomically-defined areas such as V1, V2, V3, and V4.

The action perception cluster was predominantly located in lateral parietal and premotor cortex, in regions that are classically considered as mirror-neuron system (mirror-neuron network) in humans (Cattaneo and Rizzolatti, 2009). This network, which was originally discovered in homologous regions in monkeys, is activated during action observation (Buccino et al., 2001; Caspers et al., 2010). Such activation is thought to contribute to action understanding and imitation. By testing the fMRI responses to a wide range of action categories, here we showed that this network is particularly involved in the processing of dynamic human-object interactions. Previous studies have tested the fMRI responses to static/still images of human-object interactions. These images are reportedly represented in a distributed network of areas in occipito-temporal and frontal cortex (Johnson-Frey et al., 2003; Baldassano et al., 2017). Using simple video clips of manipulative actions, studies by the Orban group have reported the involvement of the putative human anterior intraparietal sulcus (phAIP) during the observation of motor acts typically done with the hand (Jastorff et al., 2010; Abdollahi et al., 2013; Orban et al., 2019). This area was part of our action perception cluster (**Figure 2**) which showed a selective response to dynamic human-object interactions in a naturalistic setting. In our study, the responses to videos of human-object interactions were stronger in the left hemisphere, whereas the responses to videos of human-human interactions were stronger in the right hemisphere. These two lateralization effects might be complementarily associated with each other. This conjecture could be tested in an fMRI study using a large sample of subjects.

When we were investigating the time-courses of activity in the executive control clusters, we serendipitously found a large response at the transition from movie to rest. This response was not observed at the transition from rest to movie, and it was confined topographically to the executive control networks, ruling out the fact that it is merely a stimulus-driven transient signal. Since the end of each movie clip was at an unpredictable time, the large response in executive control networks could be attributed to a ‘surprise signal’. In fact, parts of the executive control network 3 were located in the cingulo-opercular cortex which plays a pivotal role in encoding surprise and salient events (Fouragnan et al., 2018). Activations in executive control networks, however, extended beyond the salience network, into regions involved in working memory (Barch et al., 2013). It is possible that, when the movie clips end abruptly, the working memory circuits are automatically activated even in the absence of an explicit cognitive task. This activation would be useful for remembering and comprehending the content and narrative of the clips. Another possibility is that the activity in executive control networks reflects disassembling an internal model of the movie events that has been progressively built up during movie-watching (Farooqui et al., 2012). This unbinding process would be more pronounced when the movie clips end.

The push-pull interaction between domain-general and domain-specific areas of cortex, reported for the first time here, has a computational benefit of using neural resources more efficiently. When the movie scenes have a clear content depicting people, objects, actions, and conversation, domain-specific areas, which are tuned to those stimuli, become active to process them. On the other hand, when the scenes are ambiguous requiring some forms of cognitive effort to resolve them, domain-general areas may be flexibly recruited. This splitting of function is in line with the idea of ‘sparse coding’ (Olshausen and Field, 2004), and it suggests that only a subset of cortical territories is active at any given point in time during movie-watching. The push-pull interaction could be tested and confirmed in subsequent studies using well-controlled paradigms. For example, by parametrically manipulating the ambiguity of naturalistic videos, one can test whether the activations shift from domain-specific to domain-general areas of cortex. The push-pull interaction could be also tested in deep neural networks by analyzing the dynamic interactions between sharply-tuned and broadly-tuned units in the top layers of networks.

What is the importance of cortical parcellation? A great deal of evidence suggests that cerebral cortex in primates is functionally compartmentalized (see Kanwisher, 2010 for review). A full characterization of the layout of cortical areas/networks would be a fundamental step in understanding the cortical computations if we assume a tight relationship between cortical organization and cortical function. Characterizing the cortical maps could also help predict what behavioral and perceptual changes would occur in pathological cases where certain regions of cortex are affected by macroscopic damage/atrophy. The cortical maps may change systematically in psychiatric disorders such as autism and schizophrenia. In future studies, cerebral cortex could be functionally parcellated in these disease populations using a movie-watching paradigm. Similarly, cerebral cortex could also be functionally parcellated in various stages of lifespan development. Such studies would shed light on how the organization of cortex changes in the course of cortical development. •

## Methods

### Subjects

In the movie-watching experiment, we used the “HCP 7T” dataset (April 2018 data release). The dataset included 184 subjects. 176 subjects (106 females, 70 males) had complete functional data for movie-watching and resting-state scans. Subjects were healthy young adults aged 22-35, and they were scanned at the Center for Magnetic Resonance Research at the University of Minnesota. The HCP data were acquired using protocols approved by the Washington University institutional review board, and written informed consent was obtained from all subjects.

In the action localizer experiment, 22 subjects (16 females, 6 males, aged 22-35) with normal or corrected-to-normal vision were scanned at the National Brain Mapping Lab in Iran. The experimental protocol was approved by an ethics committee in the Iran University of Medical Sciences (approval number: IR.IUMS.REC.1396.0465), and written informed consent was obtained from all subjects.

### Data acquisition

The HCP structural data were acquired using a customized 3 Tesla Siemens Connectom Skyra scanner with a standard Siemens 32-channel RF-receive head coil. At least one 3D T1w MPRAGE image and one 3D T2w SPACE image were collected at 0.7 mm isotropic resolution. The HCP fMRI data were acquired using a 7 Tesla Siemens Magnetom scanner with the Nova32 32-channel RF-receive head coil. Data were collected in four scan sessions using a multiband gradient-echo echo-planar imaging (EPI) sequence with the following parameters: repetition time (TR) = 1000 ms, echo time (TE) = 22.2 ms, flip angle = 45 deg, field of view (FOV) = 208 × 208 mm, matrix = 130 × 130, spatial resolution = 1.6 mm^3^, number of slices = 85, multiband factor = 5, image acceleration factor (iPAT) = 2, partial Fourier sampling = 7/8, echo spacing = 0.64 ms, bandwidth = 1924 Hz/Px. The direction of phase encoding alternated between posterior-to-anterior (PA) and anterior-to-posterior (AP) across runs. In 165 subjects, eye-tracking data were collected using an EyeLink S1000 system. 162 subjects had valid data in four runs. Of the 648 runs, 580 runs had a sampling rate of 1000 Hz, and 68 runs had a sampling rate of 500 Hz. The eye-tracking data provided horizontal and vertical gaze position and pupil size measures for each time-point.

The action localizer data were collected using a 3 Tesla Siemens Magnetom Prisma scanner with a standard Siemens 64-channel RF-receive head coil. For each subject, a whole-brain anatomical scan was acquired using a T1-weighted MPRAGE sequence (TR = 2000 ms, TE = 3.47 ms, flip angle = 7 deg, spatial resolution = 1 mm^3^, 256 sagittal slices, GRAPPA acquisition with acceleration factor of 2). The functional scans were based on a gradient-echo EPI sequence (TR = 2000 ms, TE = 30 ms, flip angle = 90 deg, spatial resolution = 3.5 mm^3^, 34 semi-axial slices, distance factor = 10%, GRAPPA acquisition with acceleration factor of 2). The slices were obtained in an even-odd interleaved order. The first 3 volumes of each run were discarded as dummy scans to allow for MR signal equilibration.

### Stimuli and experimental paradigm

In the movie-watching experiment, subjects passively viewed a series of audiovisual movie clips in four functional runs, each ~ 15 minutes in duration. Each run consisted of 4 or 5 clips. Clips varied in length from 1:03 to 4:19 min:sec. A 20-second period of rest, indicated by the word “REST” in white text on a black background, was inserted prior to the first movie clip, in between movie clips, and following the last movie clip. The first and third runs contained clips from independent films (both fiction and documentary) made freely available under Creative Commons license on Vimeo. The second and fourth runs contained clips from Hollywood films prepared by Cutting et al., 2012. The last clip of all runs was always a montage of brief (1.5 sec) videos, and it was included to facilitate test-retest and/or validation analyses. For a brief description of each clip, see Finn and Bandettini, 2021. Audio was delivered via Sensimetric earbuds, and movies were presented in a full-screen mode (size: 21.8° W × 15.7° H).

In the resting-state scans, subjects were instructed to keep their eyes open and maintain relaxed fixation on a bright cross-hair on a dark background in a darkened room. Resting-state fMRI data were acquired in four runs of approximately 16 minutes each.

In the action localizer experiment, subjects were presented with stimuli from the Action Dataset (https://data.mendeley.com/datasets/8ym35td9ft). This dataset included 300 video clips (dynamic stimuli) from 5 action categories (human-object interaction, human-human interaction, object-object interaction, human action, and object motion). Each category contained 10 subcategories (see **Supplementary Figure 3** for names of subcategories), and each subcategory contained 6 example stimuli. The duration of all clips was 4.96 seconds (124 frames at a frame rate of 25 fps). Some of the action videos were originally from the UCF101 dataset (Soomro et al., 2012), and the remaining videos were obtained from YouTube. As a control condition, 60 scrambled video clips were generated by phase-scrambling of frames of 60 action videos (12 randomly selected videos from each action category). To make smooth scrambled videos, a fixed random seed was used for phase-scrambling of all frames. For each dynamic stimulus, a static stimulus (a 4.96-second static frame/image) was also generated. The frame was typically extracted from the middle of video clips. During functional scans, the stimuli (size: 10° W × 7.5° H) were embedded in a uniform gray background. All 720 stimuli were presented to each subject in 10 runs. Each run contained 12 dynamic and 12 static stimulus blocks that were presented alternately. Each stimulus block consisted of 3 randomly ordered stimuli (videos or images) from the same action category and subcategory. At each transition between stimuli, there was one second (24 frames) of overlap in which their visual contents were gradually morphed to each other, to minimize visual transient effects. Thus, block duration was 12 seconds. A 12-second blank epoch was presented at the beginning, middle, and end of each run. Throughout the scans, subjects were instructed to continuously fixate a small fixation point at the center of screen while covertly attending to the stimuli.

Subjects viewed the stimuli on a back-projected screen (1024 × 768 pixels resolution, 60-Hz refresh rate) via a mounted mirror over the head coil. In the movie-watching experiment, the stimuli were presented using E-Prime (https://pstnet.com/products/e-prime). In the action localizer experiment, the stimuli were presented using Psychtoolbox in Matlab (http://psychtoolbox.org).

### Data analysis software

The HCP data were preprocessed using the publicly released HCP pipelines (Glasser et al., 2013). The software packages used for analysis included Connectome Workbench commandline tools (http://www.humanconnectome.org/software/connectome-workbench.html), FreeSurfer, FSL, and Matlab. Connectome Workbench ‘wb_view’ GUI was used for visualization of maps.

### Analysis of structural data

Structural images (T1w and T2w) were used for extracting subcortical gray matter structures and reconstructing cortical surfaces in each subject. Volume data were transformed from native space into MNI space using a nonlinear volume-based registration. For accurate cross-subject registration of cortical surfaces, a multimodal surface matching (MSM) algorithm (Robinson et al., 2014) was used. The MSM algorithm has two versions: ‘MSMSulc’ (non-rigid surface alignment based on folding patterns) and ‘MSMAll’ (optimized alignment of cortical areas using sulcal depth maps plus features from other modalities including myelin maps, resting-state network maps, and visuotopic connectivity maps). Data in our work were based on MSMAll registration. After surface and volume registration, cortical vertices were combined with subcortical gray matter voxels to form the standard ‘CIFTI grayordinates’ space (91,282 vertices/voxels with ~ 2 mm cortical vertex spacing and 2 mm isotropic subcortical voxels) (Glasser et al., 2016b).

### Analysis of movie-watching fMRI data

The movie-watching data were minimally preprocessed using the HCP pipelines (Glasser et al., 2013; Vu et al., 2017). Preprocessing included correction for spatial distortions due to gradient nonlinearity and b0 field inhomogeneity, fieldmap-based unwarping of EPI images, motion correction, brain-boundary-based registration of EPI to structural T1w scans, non-linear registration to MNI space, and grand-mean intensity normalization. Data from the cortical gray matter ribbon were projected onto the surface and then onto the standard grayordinates space using MSMAll registration. Data were minimally smoothed by a 2mm FWHM Gaussian kernel in the grayordinates space. Thus, smoothing was constrained to the cortical surface mesh in each hemisphere. Data were cleaned up for artifacts and structured noise using sICA+FIX. Minimal highpass filtering with a cutoff of 2000 s was also applied. The effect of this filter was similar to linear detrending of the fMRI signal. In each subject, data from four functional runs were concatenated after de-meaning.

In the clustering analysis, we used an agglomerative hierarchical clustering algorithm. The Euclidean distance was used as a distance metric, and Ward’s method was used for linkage.

### Analysis of resting-state fMRI data

The resting-state data were minimally preprocessed using the HCP pipelines (Glasser et al., 2013; Vu et al., 2017), and were projected onto the standard grayordinates space using MSMAll registration. Functional timeseries were cleaned/denoised using sICA+FIX. Minimal highpass filtering with a cutoff of 2000 s was also applied.

In the analysis in **Supplementary Figure 4**, the mean timeseries of an executive control network in each run of each subject was correlated with the timeseries of all cortical vertices in the ipsilateral hemisphere using Pearson r correlation. The resulting correlation maps were averaged across runs and across subjects after r-to-z Fisher transformation [z = artanh(r)]. The averaged correlation maps were converted back to the r maps using z-to-r transformation [r = tanh(z)].

### Analysis of action localizer data

The action localizer data were preprocessed and analyzed using FreeSurfer and FS-FAST (http://surfer.nmr.mgh.harvard.edu). For each subject, the cortical surfaces were computationally reconstructed by analyzing the anatomical MR images. The functional MR volumes were first skull-stripped using FSL’s brain extraction tool to create a mask of brain-only voxels. Then, all volumes were aligned to a reference volume at the middle time-point of each run using AFNI’s motion correction algorithm. In all runs of all subjects, the overall head motion was less than half of the voxel size. The next step of preprocessing was intensity normalization. Within the brain mask, the mean intensity of all voxels across all time-points was computed. The intensity value at each voxel at each time-point was then divided by the mean intensity and multiplied by 100. The functional volumes were rigidly co-registered to the same-subject anatomical volumes using boundary-based registration method, then they were projected onto an average cortical surface (‘fsaverage’) using spherical transformation. The functional values were spatially smoothed on the surface using a 2-D Gaussian kernel (full width at half maximum = 5 mm).

For each surface vertex, activations for different stimulus conditions were calculated using a general linear model (GLM). In this model, the timeseries of all runs within a session were concatenated, and a design matrix composed of stimulus-related task regressors and scan-related nuisance regressors was constructed. The timeseries were whitened by removing temporal autocorrelations. Task regressors were defined as boxcar functions convolved with a canonical hemodynamic response function. The head motion parameters produced during realignment were used in the GLM as nuisance regressors to account for residual effects of subjects’ movements. Additional nuisance variables included linear trends, quadratic trends, and mean confound. Prior to estimating beta values of the model, the first four time-points of each run after dummy scans were excluded to avoid inhomogeneity effects of the magnetic field. In each subject, the statistical activation maps were computed by vertex-wise t-test comparison between beta values. The group-average activation maps were obtained by mixed-effects averaging of individual subjects’ maps.

## Data/Code availability

The movie-watching and resting-state fMRI data used in this manuscript are part of the publicly available and anonymized HCP database (https://www.humanconnectome.org). The parcellation maps and related files are available for download in the BALSA database (https://balsa.wustl.edu/study/V6D4z). All analysis codes are available for sharing upon request.

## Authors’ contributions

R.R. conceived the idea and designed the analyses; R.R. collected the data; R.R., H.X., A.F., and S.K. performed the analyses and prepared the figures; R.R. wrote the manuscript; J.D. and R.D. supervised the project and critically revised the manuscript. All authors approved the final version of the manuscript.

The authors declare no competing interests.

## Acknowledgments

We thank Mojan Izadkhah and Xingjian Chu for help with data analysis, Shaghayegh Karimi and members of MRI team at the National Brain Mapping Lab in Iran for help with data collection, members of HCP team for helpful advice during data analysis, and Moataz Assem, Daniel Mitchell, and Elias Issa for helpful comments on this project. This research was supported by McGovern Institute for Brain Research, Cognitive Science and Technology Council of Iran, MRC Cognition and Brain Sciences Unit (programme SUAG/045.G101400), and a Cambridge Trust scholarship.

## Supplementary Figure Captions

**Supplementary Figure 1:**
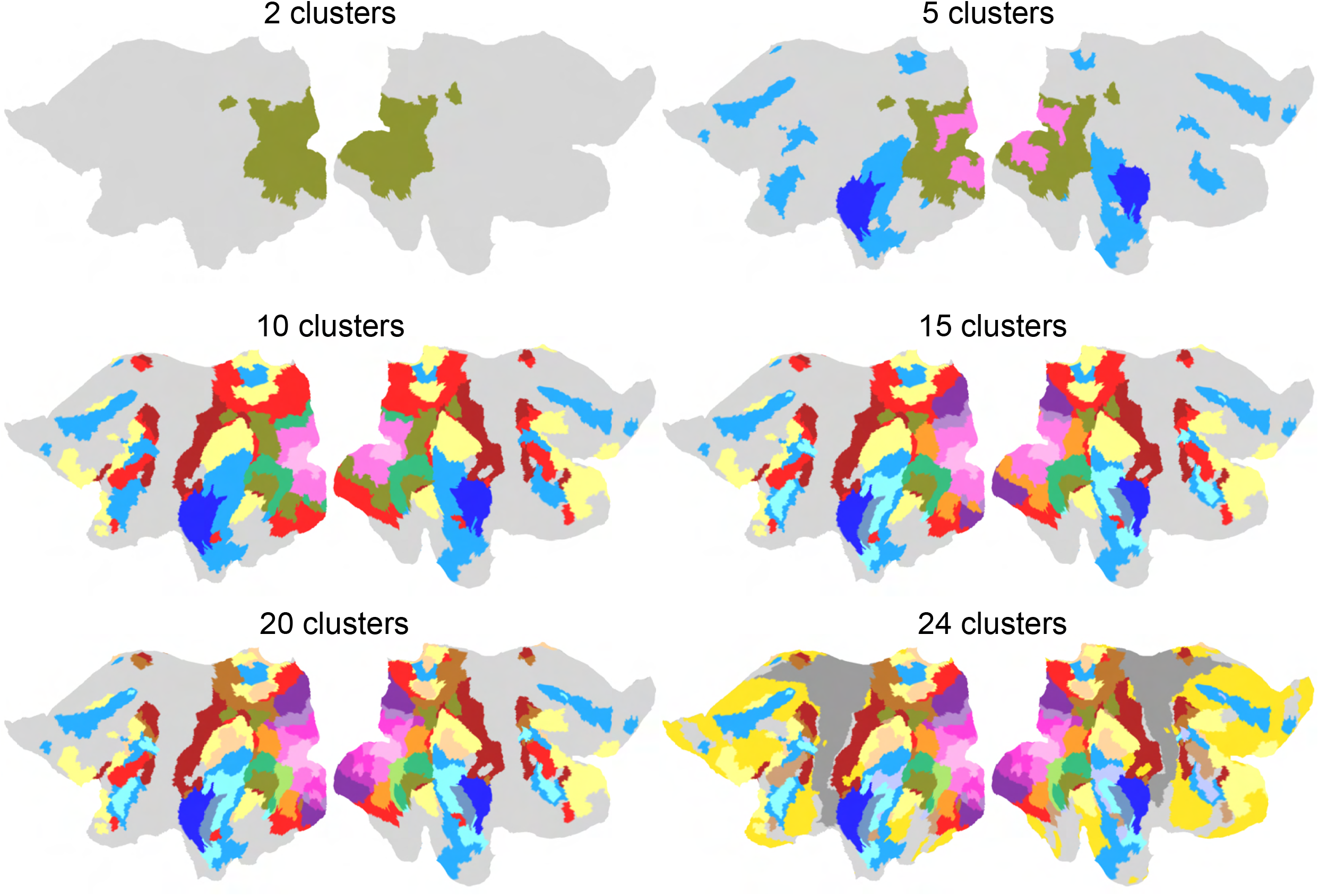
Hierarchical clustering maps. The maps are shown at the level of 2, 5, 10, 15, 20, and 24 clusters.

**Supplementary Figure 2:**
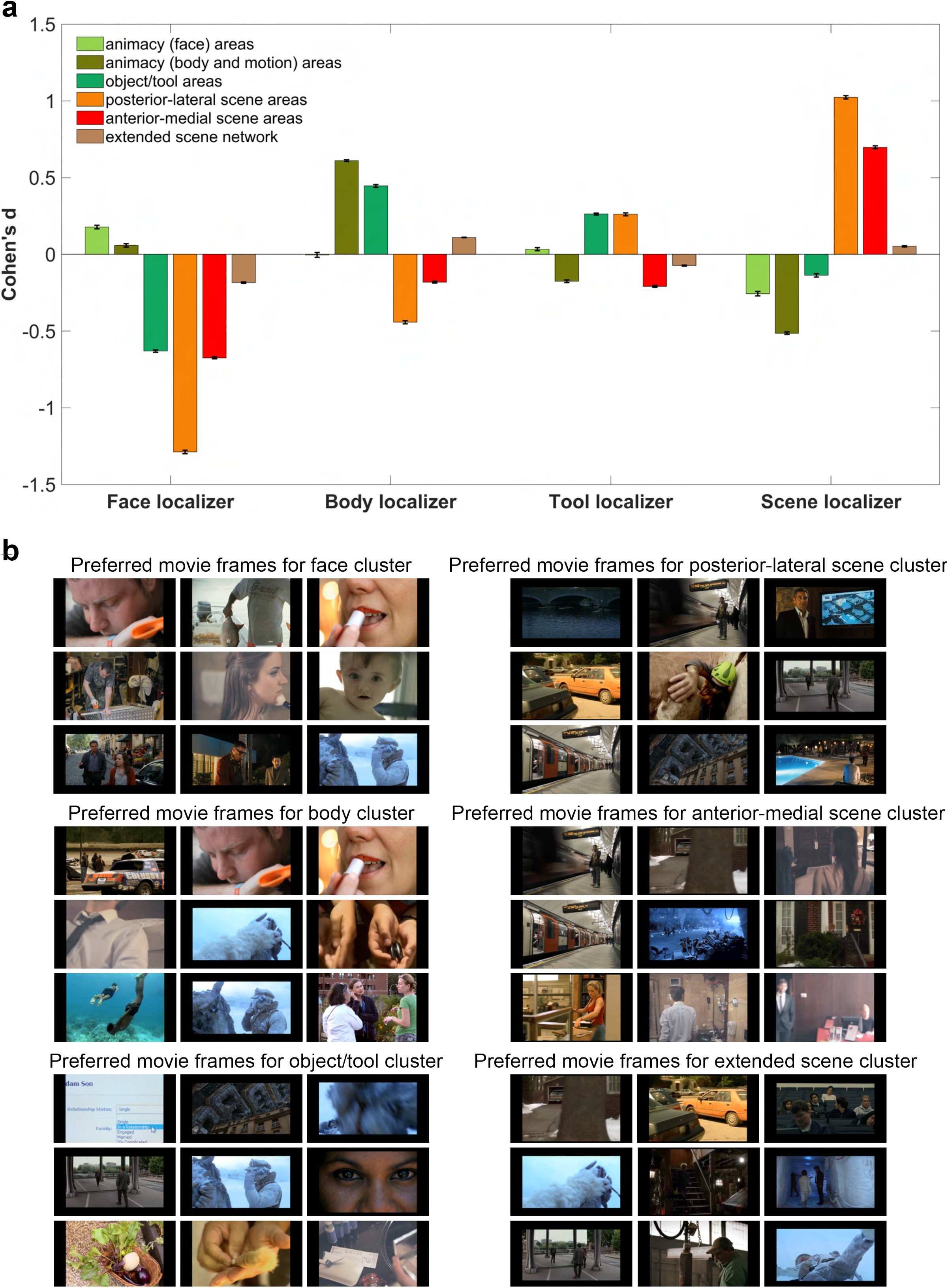
ROI analysis for category-selective clusters. (a) Using group-average data from face, body, tool, and scene localizers, an averaged activity was computed across vertices within six clusters from the movie data (animacy (face) areas, animacy (body and motion) areas, object/tool areas, posterior-lateral scene areas, anterior-medial scene areas, and extended scene network). Error bars indicate one standard error of the mean across vertices. (b) For these clusters, the first nine preferred movie frames are shown. To find the preferred movie frames of a cluster, we first obtained the mean time-course of activity across vertices of the cluster, then the peaks of response were detected using the peak-detection algorithm of Matlab. The resulting time-points were sorted based on the magnitude of response, and 50 time-points with the highest response were selected. These time-points were resorted based on an averaged activity in a 5-second window around each time-point. The movie frames corresponding to these time-points were obtained after considering a standard hemodynamic lag of 4 seconds for BOLD response (Menon et al., 1995). In the figure, the preferred movie frames are ordered consecutively first from left to right in a row then from top to bottom – where the top-left image elicited the highest response.

**Supplementary Figure 3:**
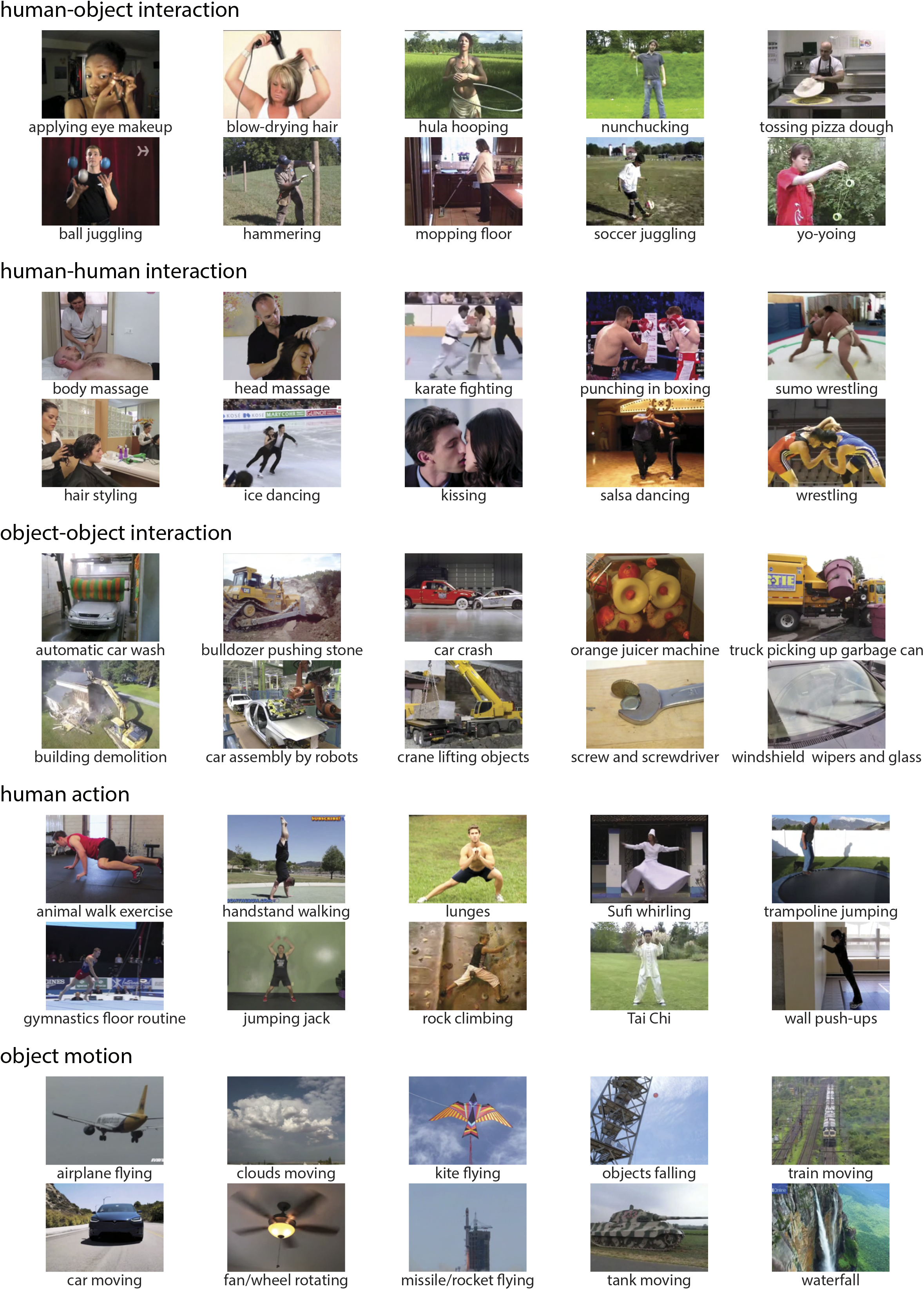
Examples of stimuli used in the action localizer experiment. The action categories included human-object interaction, human-human interaction, object-object interaction, human action, and object motion. Each category contained 10 subcategories. Each subcategory contained 6 example stimuli.

**Supplementary Figure 4:**
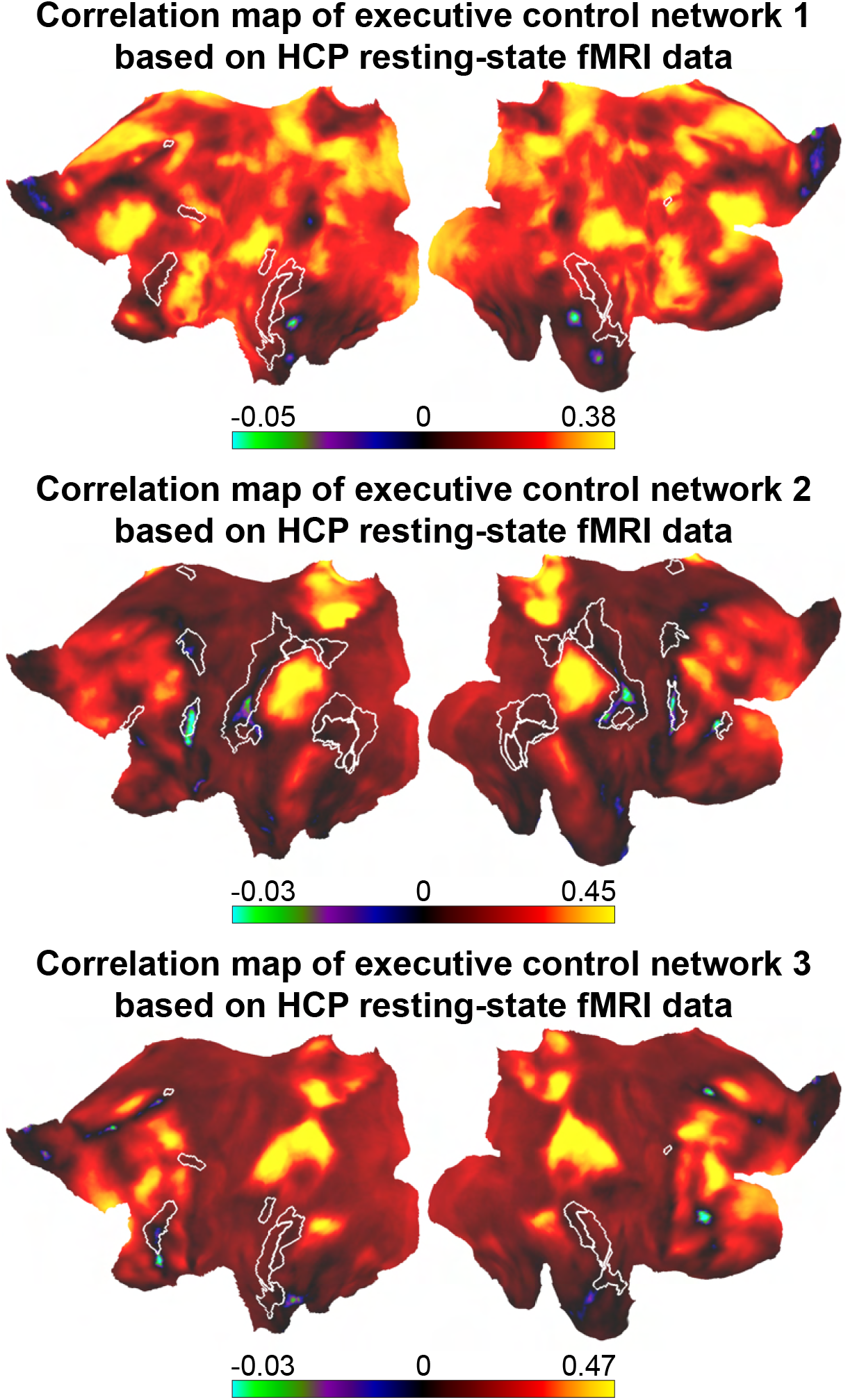
Correlation maps of executive control networks based on the HCP resting-state fMRI data. The white borders indicate domain-specific clusters shown in Figure 8a. Within these clusters, the correlation values were around zero.

**Supplementary Figure 5:**
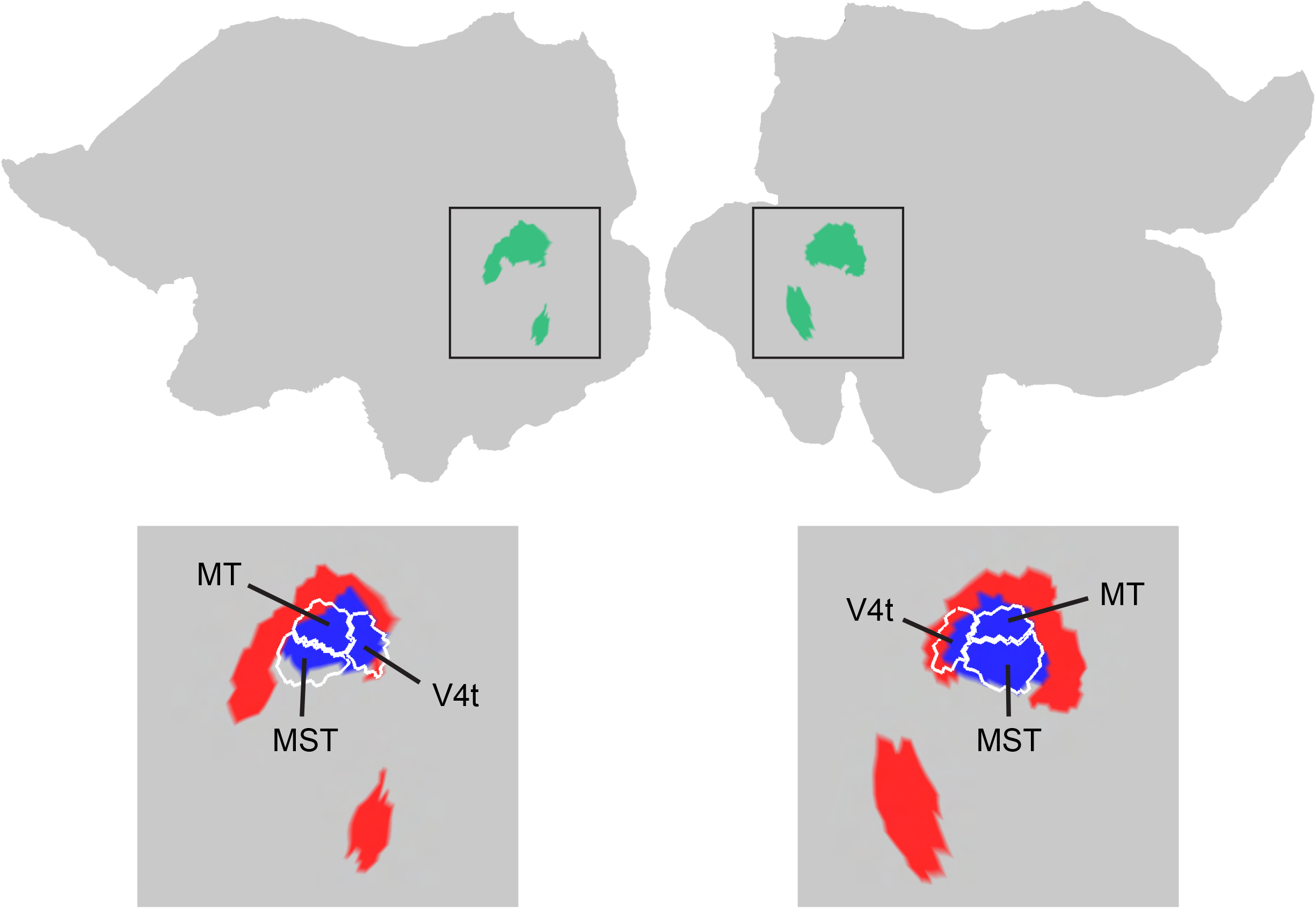
Subdivisions of the body/motion cluster. The top row shows the body/motion cluster from the parcellation map of 24 clusters. This cluster was divided into two subclusters at the level of 41 clusters (bottom row). One subcluster (blue patch) matched perfectly with MT/MST/V4t from the Glasser parcellation. The other subcluster (red patch) included EBA and FBA. As also shown previously (Weiner and Grill-Spector, 2011), EBA formed a crescent-shaped region surrounding motion areas.

## References

1. Abdollahi, R. O., Jastorff, J. & Orban, G. A.. 2013. Common and segregated processing of observed actions in human SPL. Cereb Cortex 23: 2734–53. doi: 10.1093/cercor/bhs264.

2. Amunts, K., Mohlberg, H., Bludau, S. & Zilles, K.. 2020. Julich-Brain: A 3D probabilistic atlas of the human brain's cytoarchitecture. Science 369: 988–992. doi: 10.1126/science.abb4588.

3. Amunts, K. & Zilles, K.. 2015. Architectonic Mapping of the Human Brain beyond Brodmann. Neuron 88: 1086–1107. doi: 10.1016/j.neuron.2015.12.001.

4. Arcaro, M. J., Honey, C. J., Mruczek, R. E., Kastner, S. & Hasson, U.. 2015. Widespread correlation patterns of fMRI signal across visual cortex reflect eccentricity organization. Elife 4: e03952. doi: 10.7554/eLife.03952.

5. Arcaro, M. J. & Livingstone, M. S.. 2017. A hierarchical, retinotopic proto-organization of the primate visual system at birth. Elife 6: e26196. doi: 10.7554/eLife.26196.

6. Assem, M., Glasser, M. F., Van Essen, D. C. & Duncan, J.. 2020. A Domain-General Cognitive Core Defined in Multimodally Parcellated Human Cortex. Cereb Cortex 30: 4361–4380. doi: 10.1093/cercor/bhaa023.

7. Baldassano, C., Beck, D. M. & Fei-Fei, L.. 2013. Differential connectivity within the Parahippocampal Place Area. Neuroimage 75: 228–237. doi: 10.1016/j.neuroimage.2013.02.073.

8. Baldassano, C., Beck, D. M. & Fei-Fei, L.. 2017. Human-Object Interactions Are More than the Sum of Their Parts. Cereb Cortex 27: 2276–2288. doi: 10.1093/cercor/bhw077.

9. Barch, D. M., Burgess, G. C., Harms, M. P., Petersen, S. E., Schlaggar, B. L., Corbetta, M., Glasser, M. F., Curtiss, S., Dixit, S., Feldt, C., Nolan, D., Bryant, E., Hartley, T., Footer, O., Bjork, J. M., Poldrack, R., Smith, S., Johansen-Berg, H., Snyder, A. Z. & Van Essen, D. C.. 2013. Function in the human connectome: task-fMRI and individual differences in behavior. Neuroimage 80: 169–89. doi: 10.1016/j.neuroimage.2013.05.033.

10. Beauchamp, M. S., Lee, K. E., Haxby, J. V. & Martin, A.. 2003. FMRI responses to video and point-light displays of moving humans and manipulable objects. J Cogn Neurosci 15: 991–1001. doi: 10.1162/089892903770007380.

11. Brodmann, K.. 1909. Vergleichende Lokalisationslehre der Grosshirnrinde ihren Prinzipien dargestellt auf Grund des Zellenbaues. Leipzig: Barth.

12. Buccino, G., Binkofski, F., Fink, G. R., Fadiga, L., Fogassi, L., Gallese, V., Seitz, R. J., Zilles, K., Rizzolatti, G. & Freund, H. J.. 2001. Action observation activates premotor and parietal areas in a somatotopic manner: an fMRI study. Eur J Neurosci 13: 400–404.

13. Buckner, R. L. & Yeo, B. T.. 2014. Borders, map clusters, and supra-areal organization in visual cortex. Neuroimage 93 Pt 2: 292–297. doi: 10.1016/j.neuroimage.2013.12.036.

14. Caspers, S., Zilles, K., Laird, A. R. & Eickhoff, S. B.. 2010. ALE meta-analysis of action observation and imitation in the human brain. Neuroimage 50: 1148–67. doi: 10.1016/j.neuroimage.2009.12.112.

15. Cattaneo, L. & Rizzolatti, G.. 2009. The mirror neuron system. Arch Neurol 66: 557–60. doi: 10.1001/archneurol.2009.41.

16. Cohen, A. L., Fair, D. A., Dosenbach, N. U., Miezin, F. M., Dierker, D., Van Essen, D. C., Schlaggar, B. L. & Petersen, S. E.. 2008. Defining functional areas in individual human brains using resting functional connectivity MRI. Neuroimage 41: 45–57. doi: 10.1016/j.neuroimage.2008.01.066.

17. Corbetta, M. & Shulman, G. L.. 2011. Spatial neglect and attention networks. Annu Rev Neurosci 34: 569–99. doi: 10.1146/annurev-neuro-061010-113731.

18. Cutting, J. E., Brunick, K. L. & Candan, A.. 2012. Perceiving event dynamics and parsing Hollywood films. J Exp Psychol Hum Percept Perform 38: 1476–90. doi: 10.1037/a0027737.

19. Duncan, J.. 2010. The multiple-demand (MD) system of the primate brain: mental programs for intelligent behaviour. Trends Cogn Sci 14: 172–179. doi: 10.1016/j.tics.2010.01.004.

20. Engel, S. A., Glover, G. H. & Wandell, B. A.. 1997. Retinotopic organization in human visual cortex and the spatial precision of functional MRI. Cereb Cortex 7: 181–92. doi: 10.1093/cercor/7.2.181.

21. Epstein, R. A. & Baker, C. I.. 2019. Scene Perception in the Human Brain. Annu Rev Vis Sci 5: 373–397. doi: 10.1146/annurev-vision-091718-014809.

22. Farooqui, A. A., Mitchell, D., Thompson, R. & Duncan, J.. 2012. Hierarchical organization of cognition reflected in distributed frontoparietal activity. J Neurosci 32: 17373–81. doi: 10.1523/jneurosci.0598-12.2012.

23. Felleman, D. J. & Van Essen, D. C.. 1991. Distributed hierarchical processing in the primate cerebral cortex. Cereb Cortex 1: 1–47. doi: 10.1093/cercor/1.1.1-a.

24. Finn, E. S. & Bandettini, P. A.. 2021. Movie-watching outperforms rest for functional connectivity-based prediction of behavior. Neuroimage 235: 117963. doi: 10.1016/j.neuroimage.2021.117963.

25. Fischl, B., Rajendran, N., Busa, E., Augustinack, J., Hinds, O., Yeo, B. T., Mohlberg, H., Amunts, K. & Zilles, K.. 2008. Cortical folding patterns and predicting cytoarchitecture. Cereb Cortex 18: 1973–80. doi: 10.1093/cercor/bhm225.

26. Fouragnan, E., Retzler, C. & Philiastides, M. G.. 2018. Separate neural representations of prediction error valence and surprise: Evidence from an fMRI meta-analysis. Hum Brain Mapp 39: 2887–2906. doi: 10.1002/hbm.24047.

27. Fowlkes, E. B. & Mallows, C. L.. 1983. A Method for Comparing 2 Hierarchical Clusterings. J Am Stat Assoc 78: 553–569.

28. Glasser, M. F., Coalson, T. S., Robinson, E. C., Hacker, C. D., Harwell, J., Yacoub, E., Ugurbil, K., Andersson, J., Beckmann, C. F., Jenkinson, M., Smith, S. M. & Van Essen, D. C.. 2016a. A multi-modal parcellation of human cerebral cortex. Nature 536: 171–178. doi: 10.1038/nature18933.

29. Glasser, M. F., Smith, S. M., Marcus, D. S., Andersson, J. L., Auerbach, E. J., Behrens, T. E., Coalson, T. S., Harms, M. P., Jenkinson, M., Moeller, S., Robinson, E. C., Sotiropoulos, S. N., Xu, J., Yacoub, E., Ugurbil, K. & Van Essen, D. C.. 2016b. The Human Connectome Project's neuroimaging approach. Nat Neurosci 19: 1175–87. doi: 10.1038/nn.4361.

30. Glasser, M. F., Sotiropoulos, S. N., Wilson, J. A., Coalson, T. S., Fischl, B., Andersson, J. L., Xu, J., Jbabdi, S., Webster, M., Polimeni, J. R., Van Essen, D. C. & Jenkinson, M.. 2013. The minimal preprocessing pipelines for the Human Connectome Project. Neuroimage 80: 105–24. doi: 10.1016/j.neuroimage.2013.04.127.

31. Gordon, E. M., Laumann, T. O., Adeyemo, B., Huckins, J. F., Kelley, W. M. & Petersen, S. E.. 2016. Generation and Evaluation of a Cortical Area Parcellation from Resting-State Correlations. Cereb Cortex 26: 288–303. doi: 10.1093/cercor/bhu239.

32. Goulas, A., Uylings, H. B. & Stiers, P.. 2012. Unravelling the intrinsic functional organization of the human lateral frontal cortex: a parcellation scheme based on resting state fMRI. J Neurosci 32: 10238–52. doi: 10.1523/jneurosci.5852-11.2012.

33. Hasson, U., Harel, M., Levy, I. & Malach, R.. 2003. Large-scale mirror-symmetry organization of human occipito-temporal object areas. Neuron 37: 1027–41. doi: 10.1016/s0896-6273(03)00144-2.

34. Hasson, U., Levy, I., Behrmann, M., Hendler, T. & Malach, R.. 2002. Eccentricity bias as an organizing principle for human high-order object areas. Neuron 34: 479–90. doi: 10.1016/s0896-6273(02)00662-1.

35. Hasson, U., Nir, Y., Levy, I., Fuhrmann, G. & Malach, R.. 2004. Intersubject synchronization of cortical activity during natural vision. Science 303: 1634–40. doi: 10.1126/science.1089506.

36. Haxby, J. V., Guntupalli, J. S., Connolly, A. C., Halchenko, Y. O., Conroy, B. R., Gobbini, M. I., Hanke, M. & Ramadge, P. J.. 2011. A common, high-dimensional model of the representational space in human ventral temporal cortex. Neuron 72: 404–16. doi: 10.1016/j.neuron.2011.08.026.

37. He, B. J., Snyder, A. Z., Vincent, J. L., Epstein, A., Shulman, G. L. & Corbetta, M.. 2007. Breakdown of functional connectivity in frontoparietal networks underlies behavioral deficits in spatial neglect. Neuron 53: 905–18. doi: 10.1016/j.neuron.2007.02.013.

38. Hubert, L. & Arabie, P.. 1985. Comparing partitions. J Classif 2: 193–218.

39. Huth, A. G., de Heer, W. A., Griffiths, T. L., Theunissen, F. E. & Gallant, J. L.. 2016. Natural speech reveals the semantic maps that tile human cerebral cortex. Nature 532: 453–458. doi: 10.1038/nature17637.

40. Jastorff, J., Begliomini, C., Fabbri-Destro, M., Rizzolatti, G. & Orban, G. A.. 2010. Coding observed motor acts: different organizational principles in the parietal and premotor cortex of humans. J Neurophysiol 104: 128–40. doi: 10.1152/jn.00254.2010.

41. Ji, J. L., Spronk, M., Kulkarni, K., Repovš, G., Anticevic, A. & Cole, M. W.. 2019. Mapping the human brain's cortical-subcortical functional network organization. Neuroimage 185: 35–57. doi: 10.1016/j.neuroimage.2018.10.006.

42. Johnson-Frey, S. H., Maloof, F. R., Newman-Norlund, R., Farrer, C., Inati, S. & Grafton, S. T.. 2003. Actions or hand-object interactions? Human inferior frontal cortex and action observation. Neuron 39: 1053–1058. doi: 10.1016/s0896-6273(03)00524-5.

43. Kahnt, T., Chang, L. J., Park, S. Q., Heinzle, J. & Haynes, J. D.. 2012. Connectivity-based parcellation of the human orbitofrontal cortex. J Neurosci 32: 6240–50. doi: 10.1523/jneurosci.0257-12.2012.

44. Kanwisher, N.. 2010. Functional specificity in the human brain: a window into the functional architecture of the mind. Proc Natl Acad Sci U S A 107: 11163–70. doi: 10.1073/pnas.1005062107.

45. Kanwisher, N.. 2017. The Quest for the FFA and Where It Led. J Neurosci 37: 1056–1061. doi: 10.1523/jneurosci.1706-16.2016.

46. Kanwisher, N. & Yovel, G.. 2006. The fusiform face area: a cortical region specialized for the perception of faces. Philos Trans R Soc Lond B Biol Sci 361: 2109–28. doi: 10.1098/rstb.2006.1934.

47. Kim, D., Kay, K., Shulman, G. L. & Corbetta, M.. 2018. A New Modular Brain Organization of the BOLD Signal during Natural Vision. Cereb Cortex 28: 3065–3081. doi: 10.1093/cercor/bhx175.

48. Kriegeskorte, N., Bodurka, J. & Bandettini, P.. 2008. Artifactual time-course correlations in echo-planar fMRI with implications for studies of brain function. Int J Imaging Syst Technol 18: 345–349.

49. Laumann, T. O., Gordon, E. M., Adeyemo, B., Snyder, A. Z., Joo, S. J., Chen, M. Y., Gilmore, A. W., McDermott, K. B., Nelson, S. M., Dosenbach, N. U., Schlaggar, B. L., Mumford, J. A., Poldrack, R. A. & Petersen, S. E.. 2015. Functional System and Areal Organization of a Highly Sampled Individual Human Brain. Neuron 87: 657–70. doi: 10.1016/j.neuron.2015.06.037.

50. Levy, I., Hasson, U., Avidan, G., Hendler, T. & Malach, R.. 2001. Center-periphery organization of human object areas. Nat Neurosci 4: 533–539. doi: 10.1038/87490.

51. Malach, R., Levy, I. & Hasson, U.. 2002. The topography of high-order human object areas. Trends Cogn Sci 6: 176–184. doi: 10.1016/s1364-6613(02)01870-3.

52. Menon, R. S., Ogawa, S., Hu, X., Strupp, J. P., Anderson, P. & Uğurbil, K.. 1995. BOLD based functional MRI at 4 Tesla includes a capillary bed contribution: echo-planar imaging correlates with previous optical imaging using intrinsic signals. Magn Reson Med 33: 453–459. doi: 10.1002/mrm.1910330323.

53. Nasr, S., Devaney, K. J. & Tootell, R. B.. 2013. Spatial encoding and underlying circuitry in scene-selective cortex. Neuroimage 83: 892–900. doi: 10.1016/j.neuroimage.2013.07.030.

54. Nelson, S. M., Cohen, A. L., Power, J. D., Wig, G. S., Miezin, F. M., Wheeler, M. E., Velanova, K., Donaldson, D. I., Phillips, J. S., Schlaggar, B. L. & Petersen, S. E.. 2010. A parcellation scheme for human left lateral parietal cortex. Neuron 67: 156–70. doi: 10.1016/j.neuron.2010.05.025.

55. Nishimoto, S., Huth, A. G., Bilenko, N. Y. & Gallant, J. L.. 2017. Eye movement-invariant representations in the human visual system. J Vis 17: 11. doi: 10.1167/17.1.11.

56. Olshausen, B. A. & Field, D. J.. 2004. Sparse coding of sensory inputs. Curr Opin Neurobiol 14: 481–487. doi: 10.1016/j.conb.2004.07.007.

57. Orban, G. A., Ferri, S. & Platonov, A.. 2019. The role of putative human anterior intraparietal sulcus area in observed manipulative action discrimination. Brain Behav 9: e01226. doi: 10.1002/brb3.1226.

58. Peelen, M. V. & Downing, P. E.. 2005. Selectivity for the human body in the fusiform gyrus. J Neurophysiol 93: 603–608. doi: 10.1152/jn.00513.2004.

59. Popham, S. F., Huth, A. G., Bilenko, N. Y., Deniz, F., Gao, J. S., Nunez-Elizalde, A. O. & Gallant, J. L.. 2021. Visual and linguistic semantic representations are aligned at the border of human visual cortex. Nat Neurosci 24: 1628–1636. doi: 10.1038/s41593-021-00921-6.

60. Rajimehr, R., Firoozi, A., Rafipoor, H., Abbasi, N. & Duncan, J.. 2021. Complementary hemispheric lateralization of language and social processing in the human brain. doi: 10.21203/rs.3.rs-808005/v1.

61. Rajimehr, R., Young, J. C. & Tootell, R. B.. 2009. An anterior temporal face patch in human cortex, predicted by macaque maps. Proc Natl Acad Sci U S A 106: 1995–2000. doi: 10.1073/pnas.0807304106.

62. Robinson, E. C., Jbabdi, S., Glasser, M. F., Andersson, J., Burgess, G. C., Harms, M. P., Smith, S. M., Van Essen, D. C. & Jenkinson, M.. 2014. MSM: a new flexible framework for Multimodal Surface Matching. Neuroimage 100: 414–26. doi: 10.1016/j.neuroimage.2014.05.069.

63. Russ, B. E. & Leopold, D. A.. 2015. Functional MRI mapping of dynamic visual features during natural viewing in the macaque. Neuroimage 109: 84–94. doi: 10.1016/j.neuroimage.2015.01.012.

64. Schaefer, A., Kong, R., Gordon, E. M., Laumann, T. O., Zuo, X. N., Holmes, A. J., Eickhoff, S. B. & Yeo, B. T. T.. 2018. Local-Global Parcellation of the Human Cerebral Cortex from Intrinsic Functional Connectivity MRI. Cereb Cortex 28: 3095–3114. doi: 10.1093/cercor/bhx179.

65. Soomro, K., Zamir, A. R. & Shah, M.. 2012. UCF101: A dataset of 101 human actions classes from videos in the wild. arXiv preprint arXiv:12120402.

66. Vu, A. T., Jamison, K., Glasser, M. F., Smith, S. M., Coalson, T., Moeller, S., Auerbach, E. J., Uğurbil, K. & Yacoub, E.. 2017. Tradeoffs in pushing the spatial resolution of fMRI for the 7T Human Connectome Project. Neuroimage 154: 23–32. doi: 10.1016/j.neuroimage.2016.11.049.

67. Vul, E., Lashkari, D., Hsieh, P. J., Golland, P. & Kanwisher, N.. 2012. Data-driven functional clustering reveals dominance of face, place, and body selectivity in the ventral visual pathway. J Neurophysiol 108: 2306–22. doi: 10.1152/jn.00354.2011.

68. Wang, D., Buckner, R. L., Fox, M. D., Holt, D. J., Holmes, A. J., Stoecklein, S., Langs, G., Pan, R., Qian, T., Li, K., Baker, J. T., Stufflebeam, S. M., Wang, K., Wang, X., Hong, B. & Liu, H.. 2015a. Parcellating cortical functional networks in individuals. Nat Neurosci 18: 1853–60. doi: 10.1038/nn.4164.

69. Wang, L., Mruczek, R. E., Arcaro, M. J. & Kastner, S.. 2015b. Probabilistic Maps of Visual Topography in Human Cortex. Cereb Cortex 25: 3911–31. doi: 10.1093/cercor/bhu277.

70. Weiner, K. S. & Grill-Spector, K.. 2011. Not one extrastriate body area: using anatomical landmarks, hMT+, and visual field maps to parcellate limb-selective activations in human lateral occipitotemporal cortex. Neuroimage 56: 2183–99. doi: 10.1016/j.neuroimage.2011.03.041.

71. Wig, G. S., Laumann, T. O., Cohen, A. L., Power, J. D., Nelson, S. M., Glasser, M. F., Miezin, F. M., Snyder, A. Z., Schlaggar, B. L. & Petersen, S. E.. 2014. Parcellating an individual subject's cortical and subcortical brain structures using snowball sampling of resting-state correlations. Cereb Cortex 24: 2036–54. doi: 10.1093/cercor/bht056.

72. Yeo, B. T., Krienen, F. M., Sepulcre, J., Sabuncu, M. R., Lashkari, D., Hollinshead, M., Roffman, J. L., Smoller, J. W., Zöllei, L., Polimeni, J. R., Fischl, B., Liu, H. & Buckner, R. L.. 2011. The organization of the human cerebral cortex estimated by intrinsic functional connectivity. J Neurophysiol 106: 1125–65. doi: 10.1152/jn.00338.2011.

